# Comparing alternative cholera vaccination strategies in Maela refugee camp using a transmission model

**DOI:** 10.1101/514406

**Authors:** Joshua Havumaki, Rafael Meza, Christina R Phares, Kashmira Date, Marisa C Eisenberg

**Affiliations:** Department of Epidemiology, School of Public Health, University of Michigan, 1415 Washington Heights, 48109 Ann Arbor, MI, USA; US Centers for Disease Control and Prevention; National Center for Emerging and Zoonotic Infectious Diseases; Division of Global Migration and Quarantine and Prevention, 1600 Clifton Road, 30329 Atlanta, GA, USA; US Centers for Disease Control and Prevention; Global Immunization Division - Center for Global Health, 1600 Clifton Road, 30329 Atlanta, GA, USA

**Keywords:** cholera, refugee camp, vaccination, mathematical modeling

## Abstract

**Background:** Cholera remains a major public health concern, particularly in refugee camps, which may contend with overcrowding and scarcity of resources. Maela, the largest long-standing refugee camp in Thailand, experienced four cholera outbreaks between 2005 and 2010. In 2013, a cholera vaccine campaign was implemented in the camp. To assist in the evaluation of the campaign and planning for subsequent campaigns, we developed a mathematical model of cholera in Maela.

**Methods:** We formulated a Susceptible-Infectious-Water-Recovered-based cholera transmission model and estimated parameters using incidence data from 2010. We next evaluated the reduction in cases conferred by several immunization strategies, varying timing, effectiveness, and resources (i.e., vaccine availability). Finally, we generated post-campaign case forecasts, to determine whether a booster campaign was needed.

**Results:** We found that preexposure vaccination can substantially reduce the risk of cholera even when the *<* 50% of the population is given the full two-dose series. Additionally, the preferred number of doses per person should be considered in the context of one vs. two dose effectiveness and vaccine availability. For reactive vaccination, a trade-off between timing and effectiveness was revealed, indicating that it may be beneficial to give one dose to more people rather than two doses to fewer people, given that a two-dose schedule would incur a delay in administration of the second dose. Forecasting using realistic coverage levels predicted that there was no need for a booster campaign in 2014 (consistent with our predictions, there was not a cholera epidemic in the 2014 season).

**Conclusions:** Our analyses suggest that vaccination in conjunction with ongoing water sanitation and hygiene efforts provides an effective strategy for cholera outbreaks in refugee camps. Effective preexposure vaccination depends on timing and effectiveness. If a camp is facing an outbreak, delayed distribution of vaccines can substantially alter the effectiveness of reactive vaccination, suggesting that quick distribution of vaccines may be more important than ensuring every individual receives both vaccine doses.

## Background

Global conflict, economic plight, and natural disasters interact to displace people on a large scale [1]. Over recent years, an unprecedented increase in refugee populations has led to the largest number of displaced persons ever on record [2]. Refugee camps are often built in neighboring countries to provide temporary protection and relief for refugees. However, often these crises continue for years or decades and therefore require refugees to live for an indeterminate amount of time in temporary conditions [3]. Because these refugee camps were established to provide temporary rather than permanent shelter, investments in infrastructure have not been prioritized, leading to overcrowding and poor sanitation, in turn increasing the risk of cholera and other infectious diseases [4, 5].

Cholera, a waterborne intestinal infection, causes watery diarrhea and is transmitted through fecal contamination of water and food, as well as person-to-person contact [6]. While the case fatality rate of cholera is low when treated, it can be up to 50% when untreated [7]. Approaches to reduce cholera spread include improvements in water, sanitation and hygiene (WaSH). However, political and economic hurdles can make these longer-term, larger-scale improvements challenging. Vaccination can be used as an complementary approach that results in a substantial reduction in cholera transmission [8, 9]. Further, cholera vaccines can induce herd protection, thereby reducing the risk of disease for both the vaccinated and unvaccinated segments of the population [9, 10]. Although, the effects of vaccination may attenuate over time, recent data has shown that protection remains steady several years after vaccine is administered [11]. Generally, vaccination campaigns can be implemented in conjunction with other health interventions or while more permanent preventative measures (e.g. WaSH) are being put in place.

Maela refugee camp is in northwest Thailand, 8 km east of Burma (Myanmar). In December 2009, its population was 40,009 individuals, who were mostly Burmese refugees. The distribution of ethnic groups across all Thai/Burmese border refugee camps was quite diverse, with ∼ 61% being of Karen origin [12]. Although cholera has been reported in Thailand in past years [13], a review of the literature between 1982 and 2007 found only 860 cases reported in Thailand (population of over 60 million), with the majority occurring in the northern part of the country [14]. While we do not have a denominator to determine an attack rate for these data, it is clear that compared to the outbreaks reported in the literature, the burden of cholera in Maela had been higher, with more than 1000 cases between 2005 and 2010 among a population of less than 50,000 [15]. In Maela, the majority of individuals have access water and sanitation facilities; however, sociopolitical issues and the mountainous terrain prevent its maintenance or improvement. Therefore, a vaccination campaign was implemented as a critical addition to existing efforts to reduce cholera transmission in Maela. This campaign was the first use of the oral cholera vaccine (OCV) Shanchol in a stable refugee camp [16].

An oral cholera vaccine, Shanchol, was prequalified by the World Health Organization (WHO) in 2011 [17]. The vaccine is administered in two doses, 14 days apart. Efficacy estimates from randomized control trials for the full two-dose series vary by setting and age. A large-scale age-adjusted trial conducted in India found a two-dose efficacy of 65% after follow-up at 5 years [11]. Shorter-term estimates from observational studies have found vaccine effectiveness to be as high as 86.6% after a follow up of 6 months in Guinea [18]. Notably, effectiveness of a reactive vaccination campaign in Haiti was quite close to the efficacy estimate at 63% among individuals self-reporting vaccination [19]. One-dose efficacy was estimated in a randomized trial in Bangladesh and was found to be 52% after 2 years follow up [20]. Observational studies have estimated one-dose vaccine effectiveness to be between 32.5% [21] and 87.3% [9], though the lower bound was not found to be significant. Despite logistical challenges, cholera vaccination in refugee camps has been found to be feasible and acceptable [16, 22, 23] and has been recommended as a potential key intervention to prevent and control cholera transmission in these settings.

In recent years, mathematical modeling has emerged as a useful tool in examining counterfactuals, evaluating intervention strategies, and assisting in policy decision-making [24]. Modeling of cholera transmission, in particular, has been used to guide policy and planning decisions. For example, the US Centers for Disease Control and Prevention (CDC) used real-time modeling to predict the effects of vaccination during the 2010 cholera epidemic in Haiti and to anticipate the total numbers of cases and hospitalizations [25]. More broadly, a wide range of cholera transmission models have been developed [26, 27, 28, 29, 30, 31], accounting for different mechanisms, including spatial dynamics [32, 29, 33], age structure [34, 35], environmental drivers [36, 37, 38, 39], and disease transmission characteristics such as proportion of asymptomatic individuals [38], hyperinfectiousness [40], dose response effects [41, 27, 29, 42], and multiple transmission pathways [28, 30]. One particularly relevant modeling analysis examined the impact of one compared with two doses of cholera vaccine in Haiti, Zimbabwe, and Guinea when supplies are limited [43]. Another used an agent-based model to examine cholera transmission in a refugee camp setting [44]. These models have been useful in explaining different drivers of transmission and evaluating proposed interventions. However, to our knowledge, none have explicitly considered the effects of cholera vaccination on disease transmission in a refugee camp. A wide range of characteristics such as overcrowding, high levels of population mixing, limited access to clean water and medical care, and political barriers to different interventions make it necessary to develop specific models to assess the effects of vaccination using population and outbreak data directly from refugee settings.

In this study, we the used the Susceptible-Infectious-Water-Recovered (SIWR) modeling framework [28], expanded to include two-stage vaccination and an adult/child age structure. The SIWR modeling framework accounts for both indirect transmission through environmental water sources as well as a direct pathway representing transmission through food, household water sources, and person-to-person contact. This model is an extension of the classical SIR model (with Susceptible, Infectious, and Recovered compartments to track the total number of individuals at different stages of the disease), with an additional water or environmental compartment representing the concentration of pathogen. The SIWR model has been applied to a range of cholera outbreaks as well as several theoretical studies [32, 42, 29, 28, 45]. It has been integrated with a gravity model to determine how distance and population sizes affect the spread of cholera in Haiti [32] and it was used to estimate the basic reproduction number (ℛ_0_) in a range of settings [29, 30, 32]. Finally, age-structured SIWR models [46, 34, 47] have been used to represent different transmission rates by demographic group.

The primary goal for this study was to develop a transmission model to evaluate a recent vaccine campaign in Maela refugee camp, and use this model to plan for future vaccine campaigns. The Maela refugee camp study setting is particularly relevant given the recent worldwide increase in total numbers of refugees—the approach considered here can be generalized to inform vaccine campaign planning in a wide range of contexts. Because our modeling analysis coincided with preparation and implementation of the cholera vaccination campaign in Maela, we had the opportunity to build and expand our model iteratively based on the vaccination campaign results and implementation. Our model was thus used real-time to help predict the outcome of the campaign, to determine whether any cholera outbreaks would occur in the near future, and to evaluate the necessity of administering a booster campaign for the years following the campaign. This study illustrates how mathematical modeling can be used iteratively in intervention planning to inform policy and intervention decision-making.

## Methods

### Study Cohort and Data Collection

Maela refugee camp is situated in the northwest of Thailand, 8 km east of Burma (Figure 1). In December 2010, the population was 43,645 individuals, who were mostly Burmese refugees. In total, approximately one-third of the residents were children and two-thirds were adults. As per local nongovernmental organization data collected by Premiére Urgence Aide Medicale Internationale (PU-AMI) in the camp, residents who were 15 years of age or older were considered adults because that was the working age in the camp [15]. Four cholera outbreaks occurred in Maela between 2005 and 2010. The second-largest and most recent outbreak occurred in 2010, when 362 cholera cases were confirmed by isolation of toxigenic Vibrio cholerae O1 [15].

**Figure 1.**
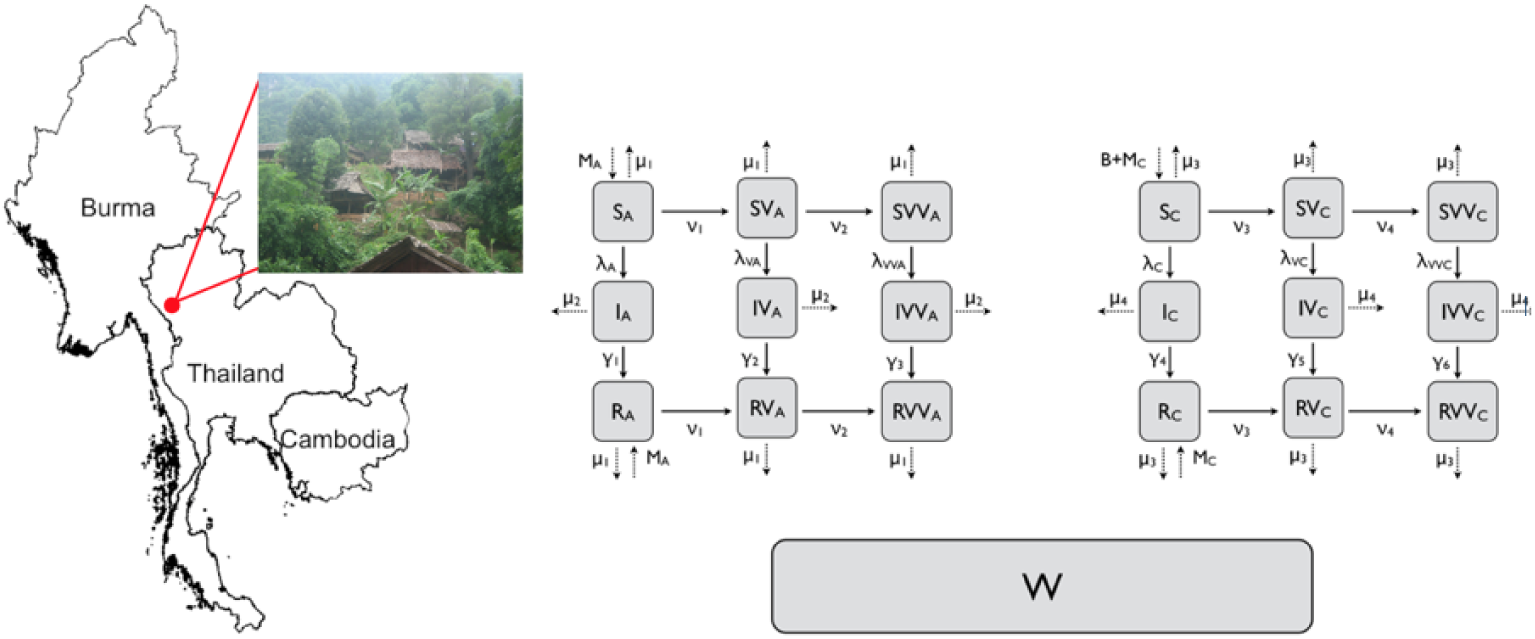
Maela Map and Model Schematic. Left: Map showing the location of Maela refugee camp. Map figure was generated in R version 3.2.4 [48] using the ‘raster’ package [49] which uses data from [50] photograph taken by JH. Right: Flow diagram of Age-structured SIWR-based model of cholera transmission in Maela. Vital dynamics and migration rates are represented by dotted lines. All infected classes (second row) shed pathogen into the common water source (*W*), which can subsequently infect susceptible individuals depending on their vaccination status.

Following WHO prequalification of Shanchol cholera vaccine in September 2011, the Thailand Ministry of Public Health sponsored a campaign for the population of Maela. The campaign was implemented in 2013 by PU-AMI with technical support from CDC. Pregnant women and children *<* 1 year old were not given vaccine (approximately 2,000 refugees were excluded in total) per the manufacturer’s recommendations [51]. Overall approximately 81% of refugees were given at least one dose of vaccine, while 64% of refugees were given two doses [15]. A baseline census was conducted and then cholera vaccine was subsequently administered. Residents of Maela have since been and will continue to be followed prospectively for new suspected cholera cases, as well as regular laboratory testing of a subset of watery diarrhea cases in the camp [15].

Our analysis considers the total population at the time of the campaign: 45,233 individuals, of whom 27,901 are adults and 17,332 are children. We fit our model to incidence data from the 2010 outbreak, the most recent outbreak at the time of the campaign, as shown in Figure 2.

**Figure 2.**
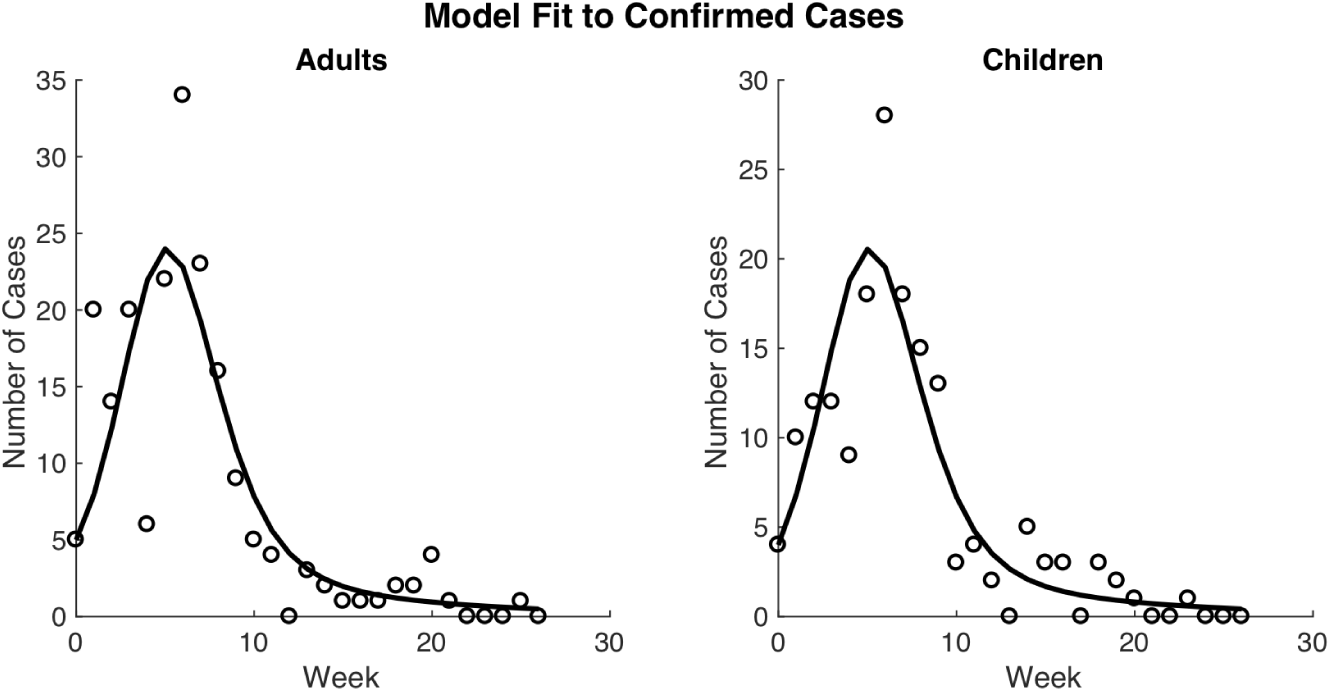
Model fit to data. Model fit to Maela cholera data from the 2010 epidemic for adults (left) and children (right). Data points for confirmed cases are indicated as circles, and simulation output is indicated as lines..

### Model Structure and Parameter Estimation

#### Age-structured SIWR Model

The model structure is shown in Figure 1. The population of Maela was separated into six model classes, non-vaccinated-adults (variables marked with *a*), once-vaccinated-adults (variables marked with *V*_*a*_), twice-vaccinated-adults (variables marked with *V V*_*a*_), non-vaccinated-children (variables marked with *c*), once-vaccinated children (variables marked with *V*_*c*_) and twice-vaccinated-children (variables marked with *V V*_*c*_). Each class is further broken into Susceptible-Infectious-Recovered compartments. Any infectious individual can infect any susceptible individual regardless of model class. The force of infection, *λ*, is the rate at which susceptible individuals become infectious (incidence rate). Children were defined as individuals *<* 15 years old. Environmental water source contamination is tracked in a separate compartment (*W*), into which any infected individual can shed. Additionally, the pathogen concentration in the water, (*W*), contributes to the calculation of *λ*.

The equations for the full model are given in the Supporting Information. Below are the force of infection equations for each demographic class and the environmental pathogen equations which are not explicitly represented in model schematic are shown in Figure 1.

#### Force of Infection Equations

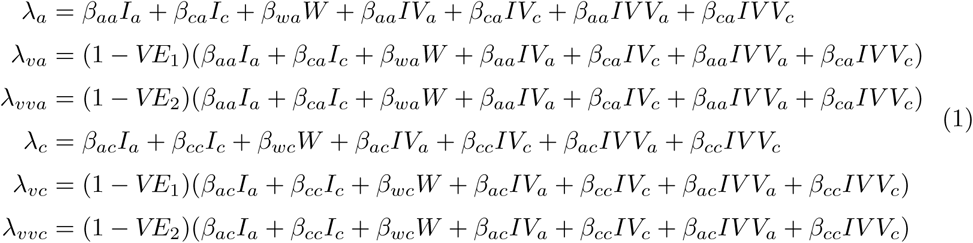

#### Environmental Pathogen

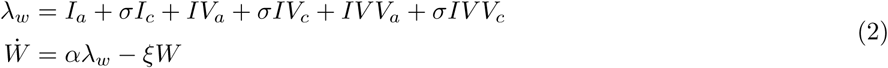

Within the force of infection equations, the *β*_*ij*_’s represent transmission from *i* to *j.*Infectious individuals shed into *W* based on *σ*, the child to adult shedding ratio of pathogen, the number of infected individuals in each class, and *α*, the rate at individuals shed into the environment. Furthermore, *ξ* is the rate at which bacteria decay in the environment. Within the remaining equations (shown in the Supporting Information and Figure 1), the *ν*’s represent the rates of vaccination for susceptible and recovered individuals. Therefore, individuals can move from nonvaccinated to once-vaccinated and then onto twice-vaccinated depending on which vaccine scenario is being simulated (see below for details on vaccination scenarios). The *µ*’s represent the combined rates for out-migration and death, and the *M*’s are the in-migration rate of susceptible or recovered individuals in Maela. *B* is the birth rate of individuals into the population. Individuals are born as susceptible children. The *γ*’s are the recovery rates of infectious individuals. All parameter definitions, values and sources are given in Table 1.

**Table 1.**
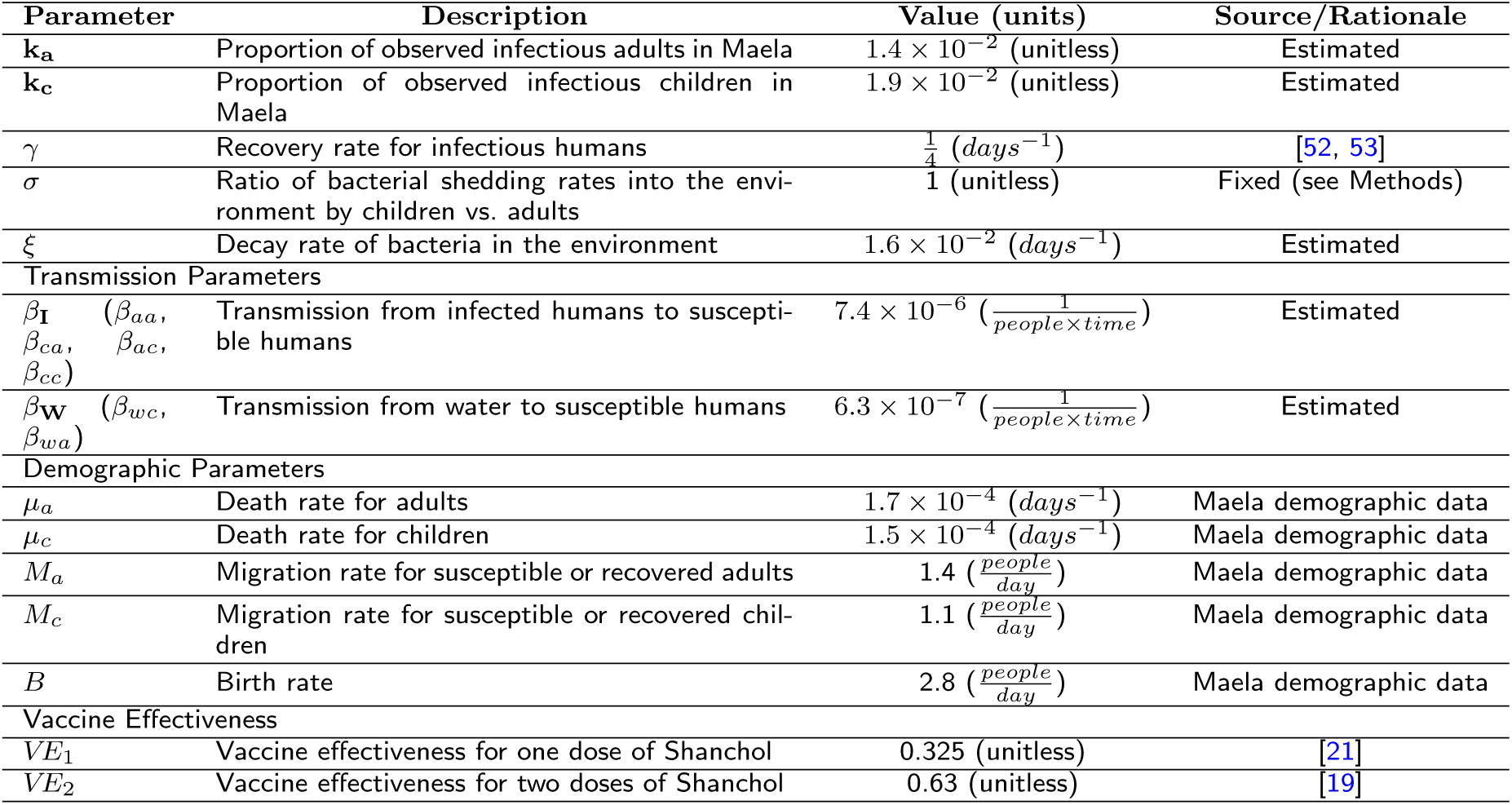
Model parameter values and estimates. Estimated parameters are indicated in bold.

#### Measurement Model

The disease surveillance data from the 2010 epidemic measure weekly cholera incidence among adults (*≥* 15 years old) and children (*<* 15 years old). To accurately represent this in the model, simulated weekly case counts are scaled by estimated reporting fraction parameters, *k*_*a*_ and *k*_*c*_. These scaling parameters represent a combination of factors including the cholera reporting rate, the fraction of asymptomatic cases (as these would not be reported), and a correction for any errors in the population size (i.e., if the population at risk is larger or smaller than the total recorded population of Maela).

#### Parameter Estimation from the 2010 Epidemic

Conducting an identifiability analysis is a necessary step before estimating parameter values from a fitted model. Identifiability is typically broken into two broad categories. First, structural identifiability is used to examine how the structure of the model and measured variables can affect what parameters are capable of being estimated assuming perfect, noise-free data [54, 30, 55, 56, 57, 58]. Once this is established, practical identifiability is used to examine the identifiability of parameter values given the actual data set being used. This step accounts for real-world data issues such as noise and sampling frequency [59, 60].We examined both structural and practical identifiability before parameter estimation. For more information on the techniques and subsequent assumptions made to reconcile identifiability issues, see Supporting Information.

We initially fit the model to estimate unknown parameters using 2010 cholera incidence data from Maela. Because the vaccine campaign was not implemented until 2013, the vaccinated compartments and relevant parameters in the full model were not included in the parameter estimation. Excluding vaccination reduced the model substantially to just three compartments for adults, three compartments for children, and the environmental pathogen compartment (*W*).

Similar to the original SIWR model [30], the structural identifiability analysis of our simplified model indicated that waterborne transmission parameters and *α* were not separately identifiable for our model, and instead formed an identifiable combination. We therefore re-scaled *W* and *β*_*W*_ to enable all unknown model parameters (*β*_*I*_, *β*_*W*_, *σ, ξ*, and the *k*’s) to be structurally identifiable (see Supporting Information for more details). From this point forward, we have only used the re-scaled versions of *β*_*W*_ and *W*. We next estimated the structurally identifiable set of unknown parameters from the data, using Poisson maximum likelihood with Nelder-Mead optimization (using `fminsearch`in Matlab).

Prior to fitting, we also fixed the demographic parameters *µ*_*a*_, *µ*_*c*_, *M*_*a*_, *M*_*c*_ and *B* based on PU-AMI data collected in the camp. The infectious period, *γ*, was fixed to 4 days [52, 53].Further, because of apparent practical identifiability limitations discovered during the initial parameter estimation (see Supporting Information for more details), we set all human-human transmission parameters denoted *β*_*I*_, equal to each other, and separately we set all human-water transmission parameters, denoted *β*_*W*_, equal to each other. We subsequently set *σ*, the child-to-adult shedding ratio of pathogen into the environment, to 1 since the *β*_*W*_’s were equal (similar to [42]). Thus the remaining parameters (*k*_*a*_, *k*_*c*_, *β*_*I*_, *β*_*W*_, *ξ*) were assumed unknown.

#### Basic Reproduction Number (ℛ_0_)

The basic reproduction number, *ℛ*_*0*_, is the number of secondary infections caused by a introduction of infectious material (individuals or pathogens) in a completely susceptible population [61]and is a commonly used measure of disease transmissibility [62]. *ℛ*_*0*_ for the model was determined using the next-generation approach [63]. For the simplified and scaled model without vaccination, *ℛ*_*0*_ is given by

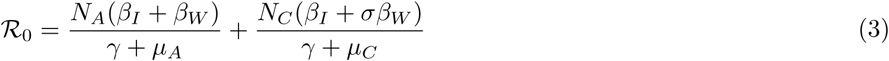

where *N*_*A*_ and *N*_*C*_ are the total adult and child populations in the camp. As we are assuming that the simulation starts near the disease free equilibrium, the total population is equivalent to the completely susceptible population (apart from the initial infected cases which begin the epidemic). Eq. (3) shows that the overall *ℛ*_*0*_ is given by a sum of the contributions by adults and children, with waterborne transmission among children weighted by the ratio of child-to-adult shedding rates (*σ*). Note that the individual terms for adult and child contributions to *ℛ*_*0*_ follow the same general form as for the original SIWR model [28].

### Vaccination Strategies: Exploration of Dynamics

To explore the effects of Shanchol on cholera transmission dynamics in this setting, we conducted hypothetical vaccination scenarios varying timing and dosage while assuming a limited amount of vaccine was administered. We incorporated different vaccination effectiveness estimates (VEs) into the transmission parameters of the vaccinated groups, and used the best-fit parameter estimates as our baseline parameters for all scenarios; see Table 1 for all parameter values. We assume that vaccination reduces the susceptibility of individuals, but that if infected, individuals are equally infectious regardless of past vaccination or cholera exposure history. We set the vaccine effectiveness in our model to be the lower bound of all estimates from recent Shanchol studies to ensure that our results do not over-estimate the impact of Shanchol. Specifically, we assumed that two doses confer an effectiveness of 63% (corresponding to the estimate from Haiti [19]) and one dose confers an effectiveness of 32.5% (corresponding to the estimate from India [21]). In all scenarios, we start with one initial observed infected individual in both *I*_*a*_ and *I*_*c*_ (i.e. 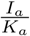 and 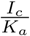, respectively) to establish consistency between scenarios and because reactive vaccination scenarios could occur only after cholera has been observed. We examine alternative pre-vaccination seeding scenarios in Supporting Information Table 5 and in the forecasting section below. All scenarios are simulated for 365 days and do not include waning immunity. Since there was only 1 death from cholera during the 2010 outbreak in Maela, we set infectious and non-infectious mortality rates equal to each other [15]. In the baseline scenario, we simulate a cholera outbreak with no vaccination campaign.

We next examined all possible one and two-dose pre-vaccination combinations (wherein all doses of vaccine are administered before the outbreak begins) by varying the proportion of the population that was given one or two-doses. As a test case for subsequent analyses, we assumed that only 20,000 doses of Shanchol were administered during the outbreak. We chose 20,000 because it was a rounded approximation of 50% of the population of Maela, which allows us to evaluate the disease dynamics in the presence of potential logistical or resource limitations. We considered two approaches: pre-vaccination and reactive vaccination (in which vaccine is administered after cholera is detected). For both approaches, we considered four general strategies:

- Two-dose: two doses of vaccine administered to 10,000 individuals. For reactive vaccination, this is implemented as two 4-day campaigns, the first of which begins one week after the first detection of cholera and the second which begins two weeks later, in accordance with Shanchol administration guidance [6].
- One-dose: single dose of vaccine administered to 20,000 individuals. For reactive vaccination, this is implemented as an 8-day campaign which begins one week after the first detection of cholera.
- Mixed: single dose of vaccine is administered to 10,000 individuals, and two doses of vaccine are administered to another 5,000 individuals.
- First come, first served: This strategy is structured similarly to the two-dose strategy, except that we do not track or control the proportion who receive one or two-doses—the 20,000 doses are administered on a first come, first served basis in two 4-day campaigns (with no prespecified number of single doses or two doses). This results in ∼ 14, 000 individuals receiving one dose and ∼ 3, 000 individuals receiving two doses.

Unless otherwise indicated, we distributed vaccine doses proportionally among adults and children. The durations of the reactive vaccine campaigns were determined based on estimates of the number of doses that could be administered per day in Maela informed by the 2013 campaign. In all reactive vaccination strategies, we assume that it takes 1 week for antibodies to confer protection. One week was chosen because a significant increase in titer levels occurred 7 days post immunization, and no significant increase in titers levels occurred beyond that time point in Shanchol seroconversion studies [64, 65]. Therefore, vaccine administered on day 7 of a cholera outbreak will not confer protection until day 14.

Given the uncertainty and variability in effectiveness estimates (they depend greatly on study design features like length of follow up), we conducted two additional analyses. First, we simultaneously varied one-dose effectiveness from 0 to 63% and number of total doses administered from 0 to twice the total camp population for all pre-vaccination scenarios. The maximum possible one-dose effectiveness was assumed to correspond to the two-dose effectiveness estimate. Second, we examined different reactive vaccination delays from 0 to 8 weeks and simulataneously varied the one-dose effectiveness from 0% to 63%.

Model simulation and parameter estimation used the `ode23tb` solver and the `fminsearch` optimization function in MATLAB [66].

### Maela Campaign Forecasting

We next used the actual campaign coverage information for Maela to forecast the potential effects of the vaccine campaign should cholera be introduced. We forecasted the total case counts that would occur in the event of an introduction of cholera in two scenarios: (1) early summer 2013 immediately after the OCV campaign had ended, and (2) 2014 assuming no cholera introduction in 2013. We simulated the 2013 scenario using OCV campaign coverage and for the counterfactual scenario of no 2013 OCV campaign. The 2014 scenario was used to assess the need for a booster campaign.

As we were simulating over a longer time than in the vaccination scenario runs, we updated our simplified model to include waning immunity. For this, we supposed that the highest level of protection (*R*_*a*_ and *R*_*c*_) comes from having recovered from a natural infection. We then supposed this immunity can wane into two compartments: *V*_2*a*_ and *V*_2*c*_, equivalent to receiving two doses of vaccine, which then wane into *V*_1*a*_ and *V*_1*c*_, equivalent to receiving only one dose of vaccine. The *V*_1*a*_ and *V*_1*c*_ compartments then wane into the susceptible classes, *S*_*a*_ and *S*_*c*_. In this model, vaccinated are considered in the appropriate *V*_2_ or *V*_1_ compartment, rather than the *SV V* and *SV* compartments, and we assume that if a vaccinated individual is infected, they simply join the overall *I*_*a*_ and *I*_*c*_ classes, so that once recovered they are fully immune (*R*_*a*_ and *R*_*c*_). For the resulting model equations, see the Supporting Information.

For initial conditions, we used the vaccination coverage data from the actual campaign in the total camp population (i.e., included and excluded individuals) to determine initial vaccination status, wherein 22.3% of adults were vaccinated with only one dose, 49.3% of adults were vaccinated with two doses, 18.7% of children were vaccinated with only one dose, and 63.6% of children were vaccinated with two doses. We note that we adjusted the percent coverage of individuals included in the campaign from that reported in [15] to account for the total camp coverage of both included and excluded individuals. See the Supporting Information for details on these calculations. In addition to the vaccination campaign, we also considered the possibility of pre-existing immunity (because there had been cholera epidemics previously in the camp). We therefore ran all scenarios with two options: assuming a fully susceptible population, and alternatively a partially immune population (up to 50%). We denote the fraction of the initial population which is immune to be *F*_*imm*_ (split evenly across the immune compartments). We also allowed the fraction of immune individuals (assumed to all be at V 1-level immunity) in incoming migrations, denoted *M*_*imm*_, to range from 0 to 1. Finally, to model the introduction of cholera, we added a single adult case and a single child case into the model. For a cholera introduction in early summer 2013, each of the following scenarios was run:

- No OCV campaign in a fully susceptible population
- OCV campaign in a fully susceptible population
- No OCV campaign in a partially immune population
- OCV campaign in a partially immune population

For a cholera introduction in 2014, we ran the model without cholera introduction for 2013 (to allow the waning immunity, migration, birth, and death dynamics to continue), and then for 2014 each of the following scenarios was run twice (once seeding with observed cases and once seeding with actual cases):

- 2013 OCV campaign in a fully susceptible population
- 2013 OCV campaign in a partially immune population

To incorporate uncertainty in our parameter values into our projections, we used Latin hyper-cube sampling (LHS). We sampled 1,000 parameter sets for each forecasting scenario using the parameter ranges shown in Table 2. For most parameters, we based our ranges on the maximum and minimum values observed from the Maela demographic data from 2009 to 2013, as well as parameter ranges evaluated from the literature (see Table 2). Although the ranges of one compared with two-dose effectiveness overlap in studies, we assume that the maximum one-dose effectiveness is equal to the minimum two-dose effectiveness. For the remaining transmission and reporting rate parameters, we used broad *±* 50% ranges. The ranges used for the estimated parameters were generally similar to or wider than their estimated confidence bounds but may better reflect the additional uncertainties coming from the fact that each introduction of cholera may be different (in time of year, contact patterns among the population, reporting, ongoing interventions, etc.).

**Table 2.** LHS ranges among fitted parameters.

| Parameter | Range | Source |
| --- | --- | --- |
| $\beta_I$ | $0.5 \times \beta_I$ to $1.5 \times \beta_I$ | See Table 1 for best-fit $\beta_I$ used |
| $\beta_W$ | $0.5 \times \beta_W$ to $1.5 \times \beta_W$ | See Table 1 for best-fit $\beta_W$ used |
| $k_a$ | $0.5 \times k_a$ to $1.5 \times k_a$ | See Table 1 for best-fit $k_a$ used |
| $k_c$ | $0.5 \times k_c$ to $1.5 \times k_c$ | See Table 1 for best-fit $k_c$ used |
| $\mu_a$ | 1.48e-5 to 0.0006 | Maela demographic data 2009-2013 |
| $\mu_c$ | 7.82e-6 to 0.0027 | Maela demographic data 2009-2013 |
| $M_a$ | 0.574 to 8.525 | Maela demographic data 2009-2013 |
| $M_c$ | 0.475 to 4.656 | Maela demographic data 2009-2013 |
| $B$ | 1.9 to 3.87 | Maela demographic data 2009-2013 |
| $\gamma$ | 0.2 to 0.3 | [32, 30, 39, 28, 52, 67] |
| $\alpha$ | $\frac{1}{5 \times 365}$ to $\frac{1}{2 \times 365}$ | [42, 68, 38, 69] |
| $\xi$ | $\frac{1}{365}$ to $\frac{1}{5}$ | [32, 30, 39, 28, 70, 71, 72, 73] |
| $VE_1$ | 0 to 0.67 | [74, 9] |
| $VE_2$ | 0.67 to 0.8 | [75, 15] |
| $M_{imm}$ | 0 to 1 | See Section Maela Campaign Forecasting |
| $F_{imm}$ | 0 to 0.5 | See Section Maela Campaign Forecasting |

## Results

### Parameter Estimation and Uncertainty

Figure 2 shows the model fitted to weekly incidence data for adults and children, with parameter estimates given in Table 1. To formally examine parameter uncertainty and practical identifiability, we plotted profile likelihoods for fitted parameters *k*_*a*_, *k*_*c*_, *β*_*W*_, *β*_*I*_, *ξ*, shown in Supporting Information Figure 9. All parameters were shown to be identifiable (finite confidence bounds), with clear minima in each case, although the uncertainty was comparatively high for *β*_*W*_ and *ξ*, consistent with previous studies showing that these two parameters are often practically unidentifiable [30, 42].

### Vaccination Strategies: Exploration of Dynamics

We assessed different roll-out vaccination campaign strategies to determine the most effective method of preventing a cholera outbreak in Maela.

#### Baseline Scenario – No Vaccination

The baseline scenario in which no vaccination is given yields a total of 395.4 cases with an attack rate of 8.7 per 1,000 people. See Table 3 for case counts and attack rates from the theoretical exploration of dynamics analysis.

**Table 3.** Attack rates and numbers of cases for each theoretical scenario (assuming 20,000 doses) using best fit parameter values

| Vaccination Scenario | Cases | Attack Rate (per 1,000 people) |
| --- | --- | --- |
| Baseline: No vaccination | 395.4 | 8.7 |
| Pre-Vaccination |  |  |
| One-dose | 255.3 | 5.6 |
| Two-dose | 247.6 | 5.5 |
| Mixed | 251.4 | 5.6 |
| First come, first served | 252.8 | 5.6 |
| Reactive Vaccination |  |  |
| One-dose | 267.6 | 5.9 |
| Two-dose | 268.2 | 5.9 |
| Mixed | 271.5 | 6.0 |
| First come, first served | 273.7 | 6.1 |

#### Pre-Vaccination Scenarios

We first examined how variation in pre-vaccination coverage of one compared with two doses affects cumulative case counts. As one-dose or two-dose coverages increase, case counts decrease. The cutoff for one-dose coverage alone to result in less than 50 total cases is 96% and the corresponding cutoff for two-dose coverage alone is 49%. All the pre-vaccination scenarios we considered administering 20,000 doses result in similar cumulative case counts, while the real-world Maela scenario (with higher coverage) was more effective. See Figure 3 for details.

**Figure 3.**
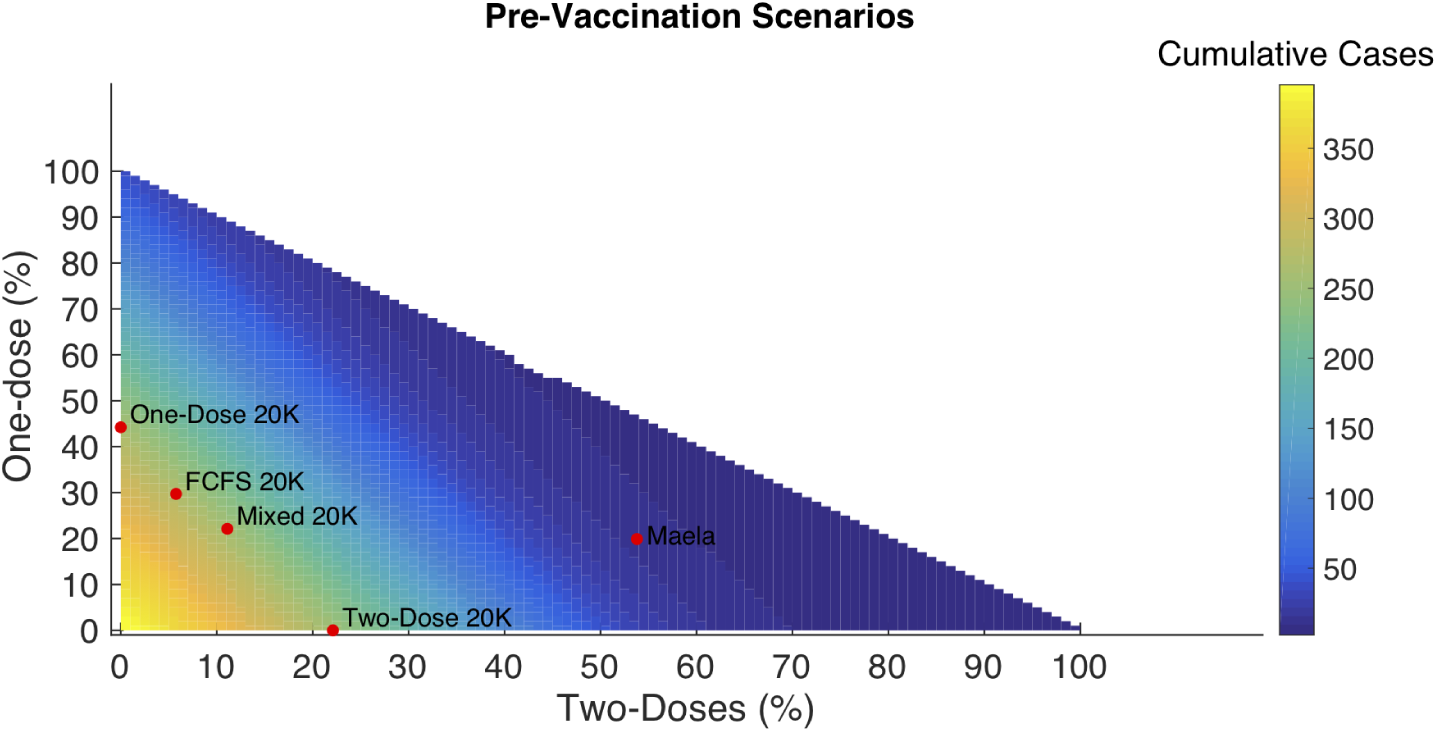
Cumulative cases varying pre-vaccination coverage. Cumulative cases resulting from methodically varying one-dose vs. two-dose coverage for pre-vaccination. With labels indicating pre-vaccination scenarios with 20,000 doses. (FCFS = first-come, first-served).

We then simulated pre-vaccination scenarios administering 20,000 doses to Maela and using the one-dose, two-dose, mixed, and first come, first served strategies. All four prevaccination campaigns yielded similar results. An outbreak occurs, but it is substantially smaller than the baseline scenario. Total case counts range from 247.6 with an attack rate of 5.6 per 1,000 people in the two-dose scenario to 255.3 with an attack rate of 5.6 per 1,000 people in the one-dose scenario. See Figure 4 and Table 3 for details.

**Figure 4.**
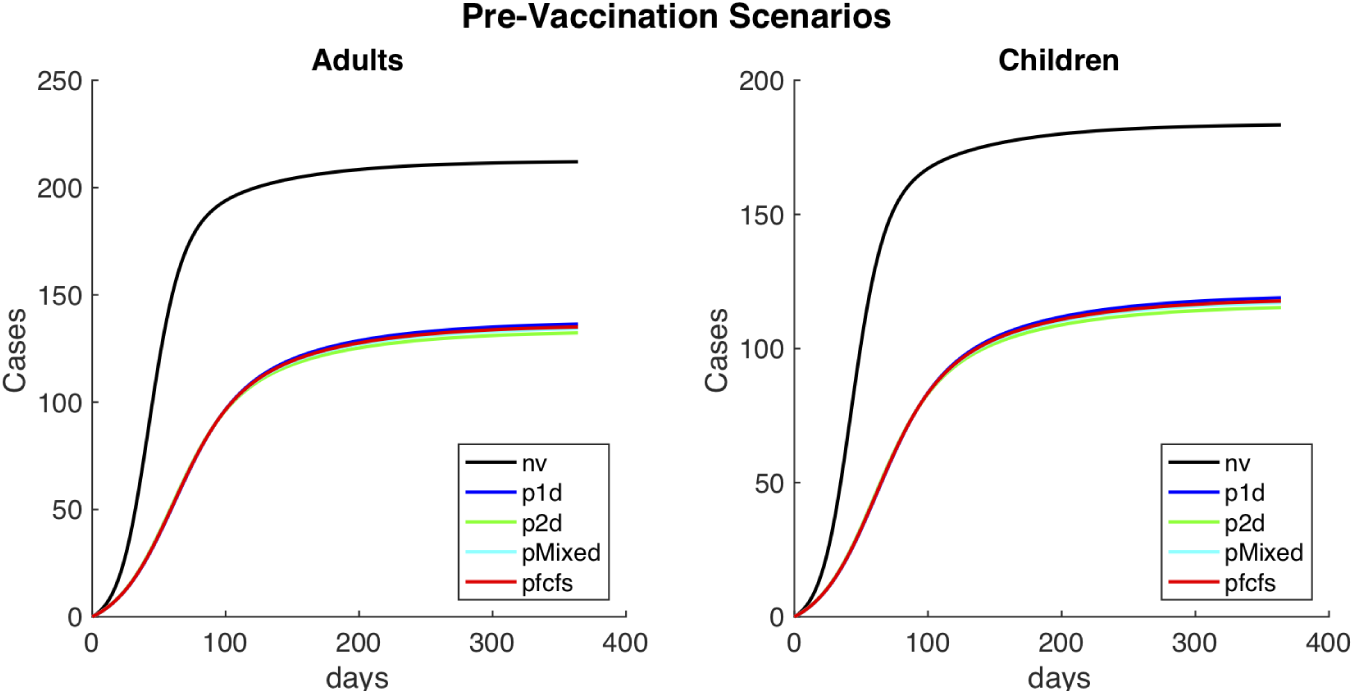
Cumulative Cases for alternative pre-vaccination scenarios. Cumulative cholera cases in adults and children, for alternative pre-vaccination scenarios. Where ‘nv’ is the baseline, no vaccination scenario; ‘p1d’ the one-dose scenario; ‘p2d’ is the two-dose scenario; ‘pmixed’ is the mixed scenario, and pfcfs is the first-come first-served scenario.

Finally, we varied one-dose effectiveness (from 0 to the full two-dose effectiveness) and total number of doses (from 0 to twice the total camp population (full coverage with two doses)) in all pre-vaccination scenarios. Intuitively, when one-dose effectiveness is low, the two-dose scenario is most effective. For instance, administering the full two-dose series to 22,000 people is sufficient to achieve *<* 50 cases. However, the one-dose strategy alone can achieve large reductions in case counts if the combination of effectiveness and number of doses is sufficient. For instance, either an effectiveness of 31% combined with 45,000 doses or an effectiveness of 63% combined with 22,000 doses are minimally sufficient to achieve *<* 50 cumulative cases. If one-dose effectiveness is very high and approximately equal to two-dose effectiveness, the two scenarios behave similarly. See Figure 5 for details.

**Figure 5.**
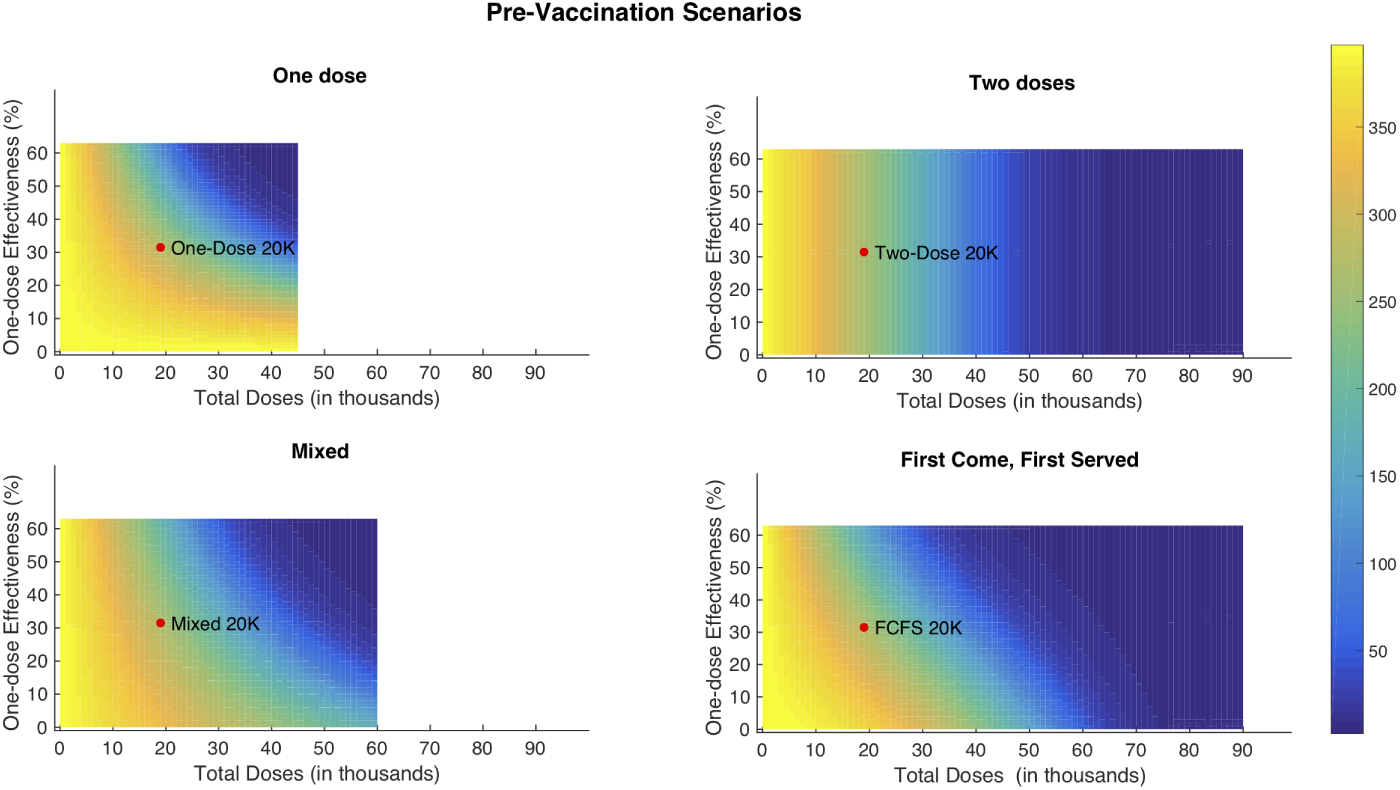
Cumulative cases varying one-dose vaccine effectiveness and total doses for different pre-vaccination scenarios. Cumulative cases resulting from methodically varying one-dose vaccine effectiveness and total doses administered in campaign implementation for the all pre-vaccination scenarios. (FCFS = first-come, first-served).

#### Reactive Vaccination Scenarios

Next, we examined reactive vaccination scenarios, again administering 20,000 doses given in separate campaigns starting 7 days after cholera is initially detected. The same one-dose, two-dose, mixed, and first come, first served scenarios were simulated, shown in Figure 6 and Table 3.

**Figure 6.**
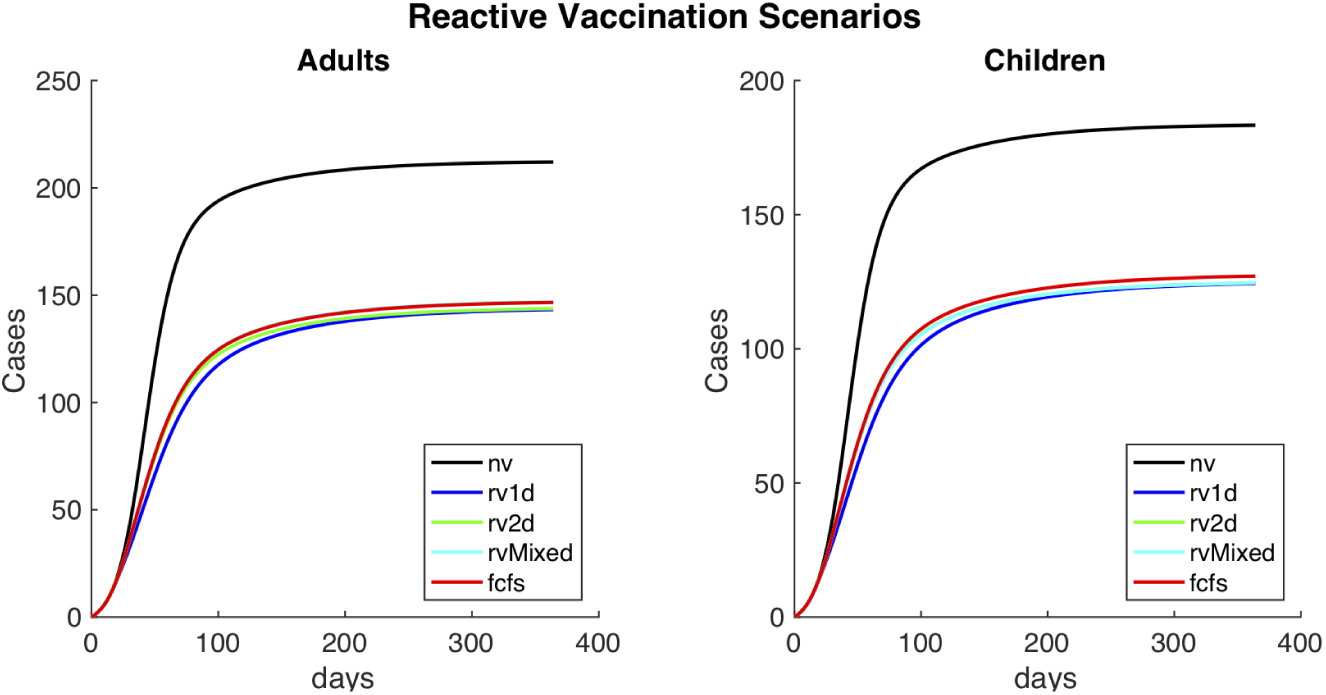
Cumulative Cases for alternative reactive vaccination scenarios. Cumulative cholera cases in adults and children, for alternative reactive vaccination scenarios. Where ‘nv’ is the baseline, no vaccination scenario; ‘rv1d’ is the one-dose scenario; ‘rv2d’ is the two-dose scenario; ‘rvMixed’ is the mixed scenario; and ‘fcfs’ is the first come, first served scenario.

In all vaccination test-case scenarios (shown in Figure 6), an outbreak occurs but it has substantially lower numbers of cases than the baseline scenario. The one-dose, two-dose, mixed, and first come, first served scenarios all yield similar results, with total case counts ranging from 267.6 with an attack rate of 5.9 per 1,000 people in the one-dose scenario to 273.7 with an attack rate of 6.1 per 1,000 people in the first come, first served scenario. Given the fact that the one-dose effectiveness is almost exactly half of the two-dose effectiveness, the one-dose scenario provides approximately the same amount of coverage as the other scenarios.

The case counts are higher and the differences between scenarios are lower in the reactive vaccination scenarios than in the pre-vaccination scenarios, indicating that pre-vaccination is more effective at reducing the spread of cholera.

Finally, to evaluate the effects of vaccine campaign timing, we methodically varied one-dose vaccine effectiveness from 0% to 63% and initial vaccination campaign onset from 0 days to 55 days (or 8 weeks) after cholera was first identified for all reactive vaccination scenarios. Results are shown in Figure 7. As with the pre-vaccination scenarios, when one-dose effectiveness is low or uncertain, the two-dose strategy is preferable. For instance, if one-dose effectiveness is 20%, the minimum number of cases (i.e. assuming no delay) for the one-dose strategy is 320 while the minimum for the two-dose strategy is 291.8. Once one-dose effectiveness is sufficiently high, all campaigns are comparable given relatively short delays. This is illustrated by the 20,000 dose reactive vaccination scenarios (see Figure 6). A one-dose effectiveness of 27% is minimally sufficient for it to be the preferable strategy across all campaigns and given any delay. When reaction time is important, one-dose is often the preferred strategy because all doses of vaccine are administered in a single 8 day campaign instead of across two 4 day campaigns separated by 14 days. This results in more population level protection earlier in the epidemic. Lastly, beyond a certain point in the epidemic, one-dose effectiveness has little impact on total case counts across all campaigns. This interplay between vaccination effectiveness and timing highlights the importance of minimizing delays in reactive vaccination campaigns with pre-vaccination being the most effective way to reduce the spread of cholera (see Figure 7).

**Figure 7.**
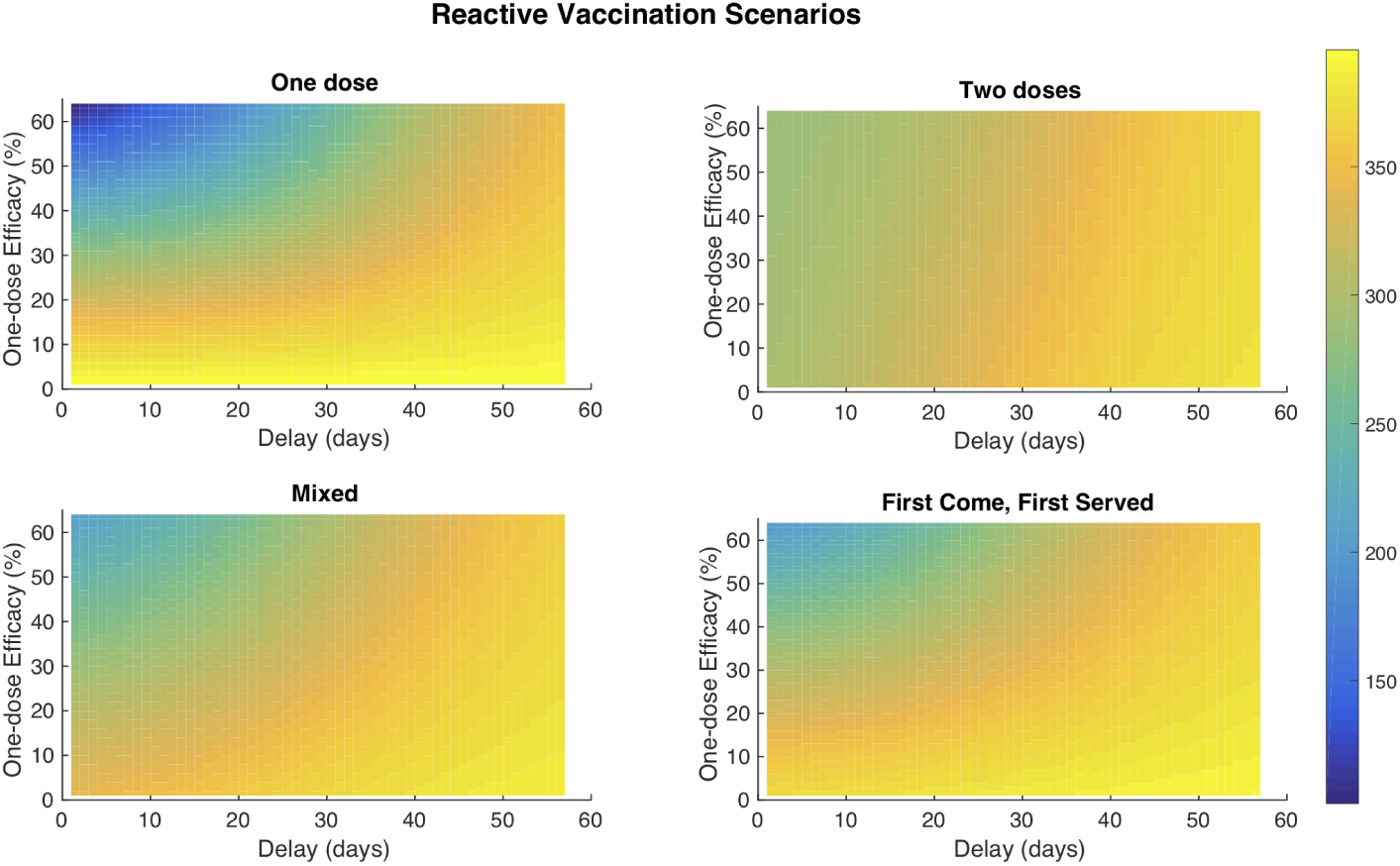
Cumulative cases varying one-dose vaccine effectiveness and vaccine campaign delay for reactive vaccination scenarios. Cumulative cases resulting from methodically varying one-dose vaccine effectiveness and delay in campaign implementation for all reactive vaccination scenarios.

### Maela Campaign Forecasting

Table 4 shows the forecasted numbers of cases and attack rate for each of the 2013 and 2014 scenarios using the parameter estimates in Table 1.

**Table 4.** Forecasts for the 2013 and 2014 cholera seasons.

| Scenario | Median Total Cases | (25%, 75% quantiles) |
| --- | --- | --- |
| 2013 Forecasts - Fully Susceptible Population |  |  |
| No OCV | 333.3 | (15.1, 540) |
| With OCV | 0.2 | (0.1, 11) |
| 2013 Forecasts - Partially Immune Population |  |  |
| No OCV | 139.9 | (0.3, 364.9) |
| With OCV | 0.1 | (0.1, 0.4) |
| 2014 Forecasts |  |  |
| With OCV - Fully Susceptible Pop. | 0.4 | (0.1, 158.8) |
| With OCV - Partially Immune Pop. | 0.3 | (0.1, 101) |

**Table 5.**
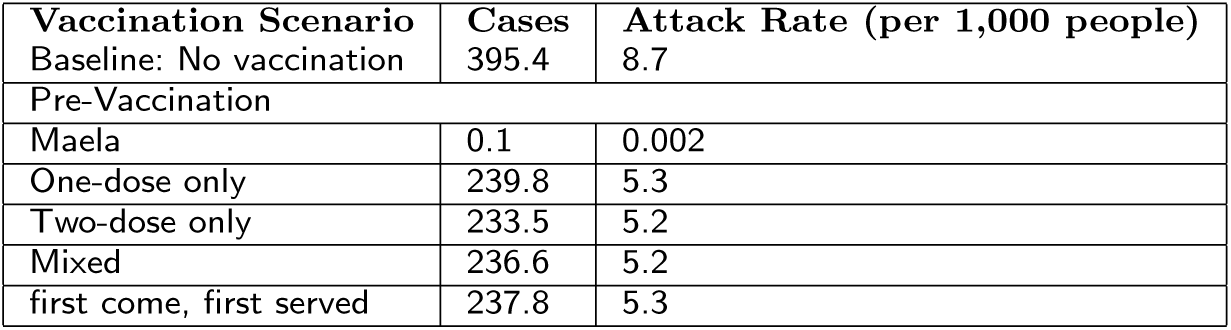
Cumulative Cases of Alternative Seeding Scenarios.

#### Forecasts for the 2013 OCV Campaign

In the 2013 forecasting results, we see a larger spread of total case numbers for runs in the scenario without the OCV campaign compared to the scenario with the OCV campaign. Case counts range from 0 to approximately 1,000 in the fully susceptible population. The partially immune population runs generally have lower case counts. Furthermore, for the scenarios that consider the OCV campaign, we see the vast majority of runs having case counts close to 0. For forecasts among a fully susceptible population, see Figure 8, and for forecasts among a partially immune population, see Supporting Information Figure 11.

**Figure 8.**
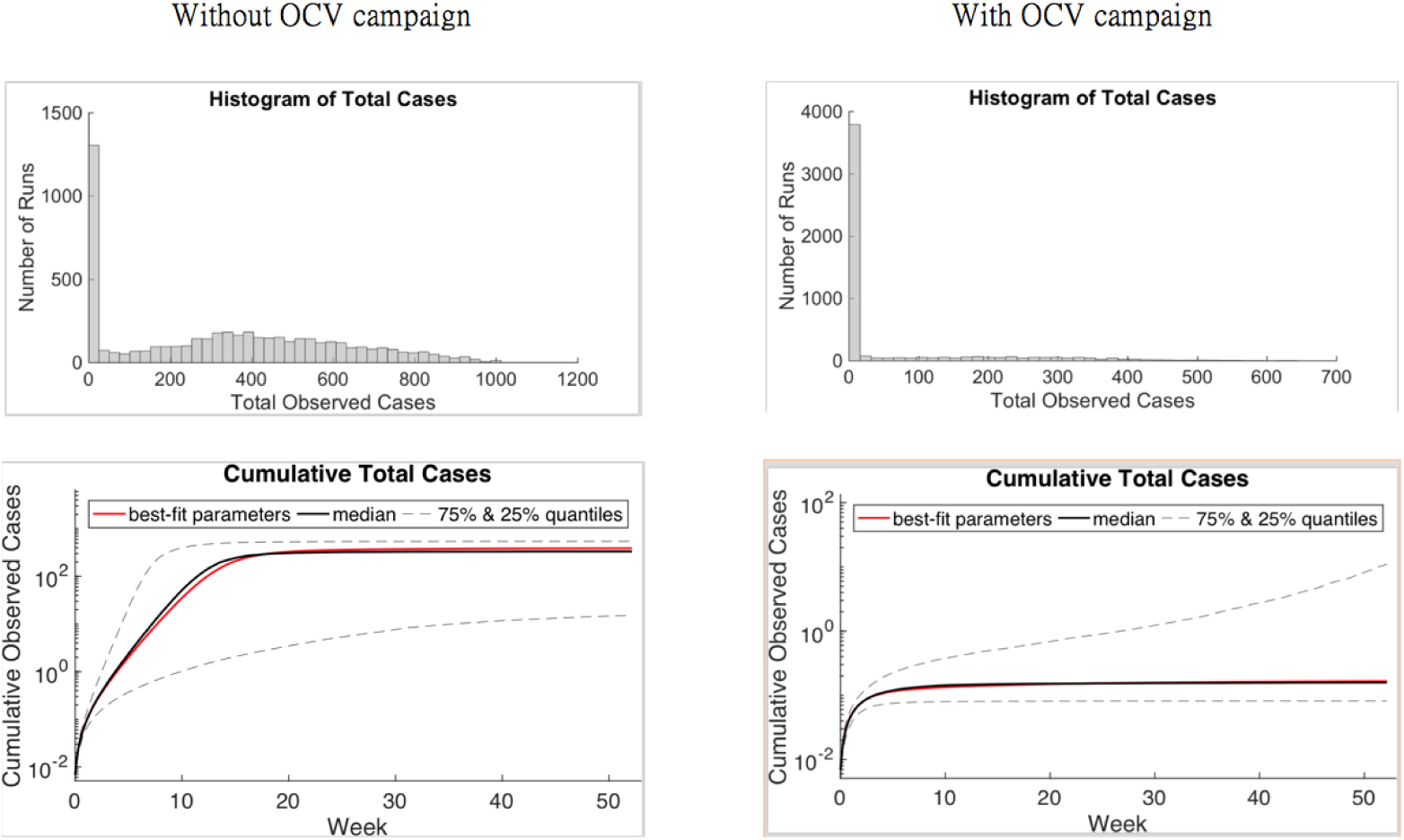
2013 forecasting results. 2013 forecast with a single actual case in adults and children as seeding. The fully susceptible population with no OCV campaign on left and with OCV campaign on right.

**Figure 9.**
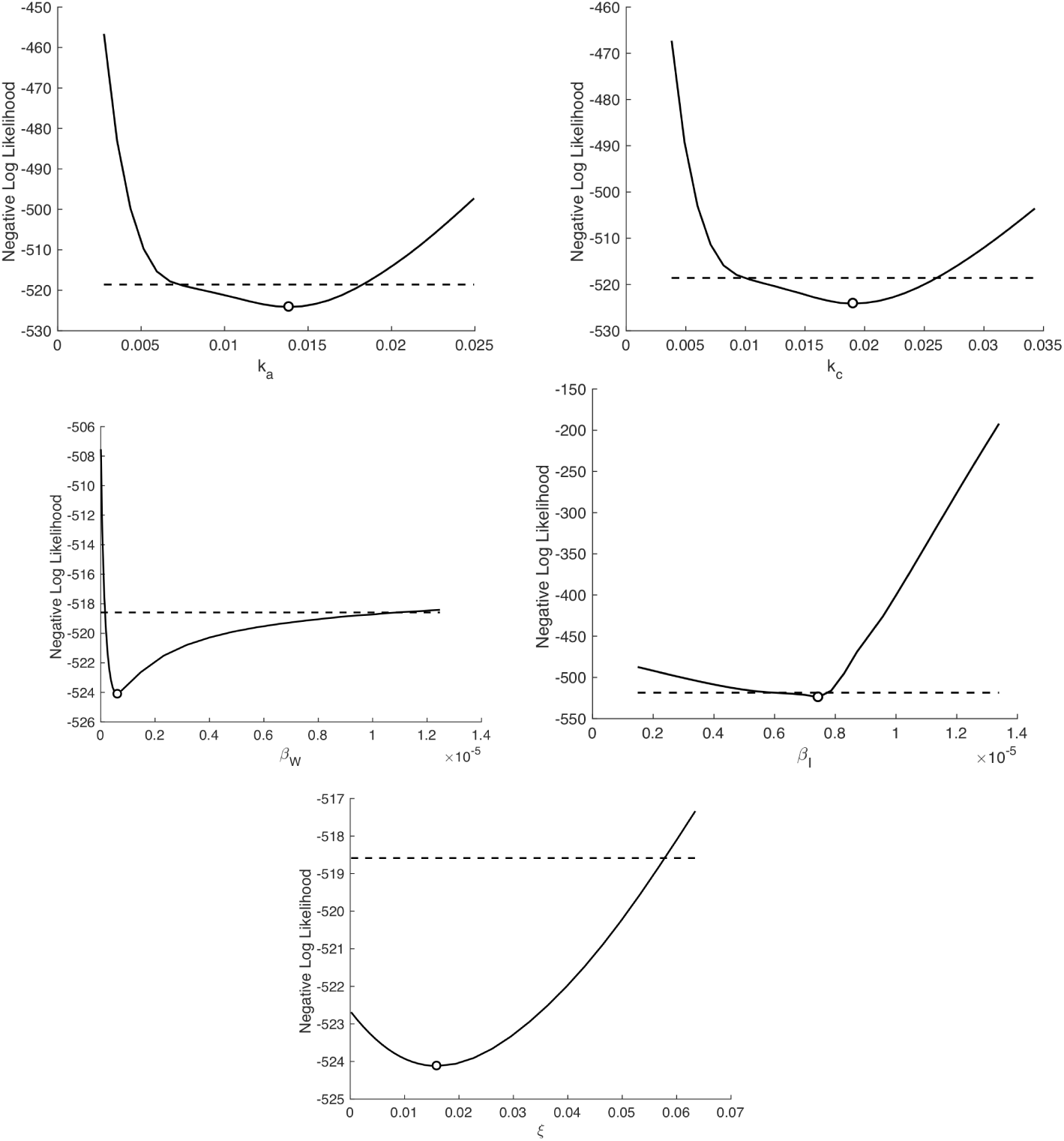
Profile likelihood plots for estimated parameters top row: *k*_*a*_ (left), *k*_*c*_ (right) second row: *β*_*W*_ (left), *β*_*I*_, and third row: *ξ* Note: The *β*_*W*_ *and ξ* ranges were extended to capture the 95 % confidence intervals.

**Figure 10.**
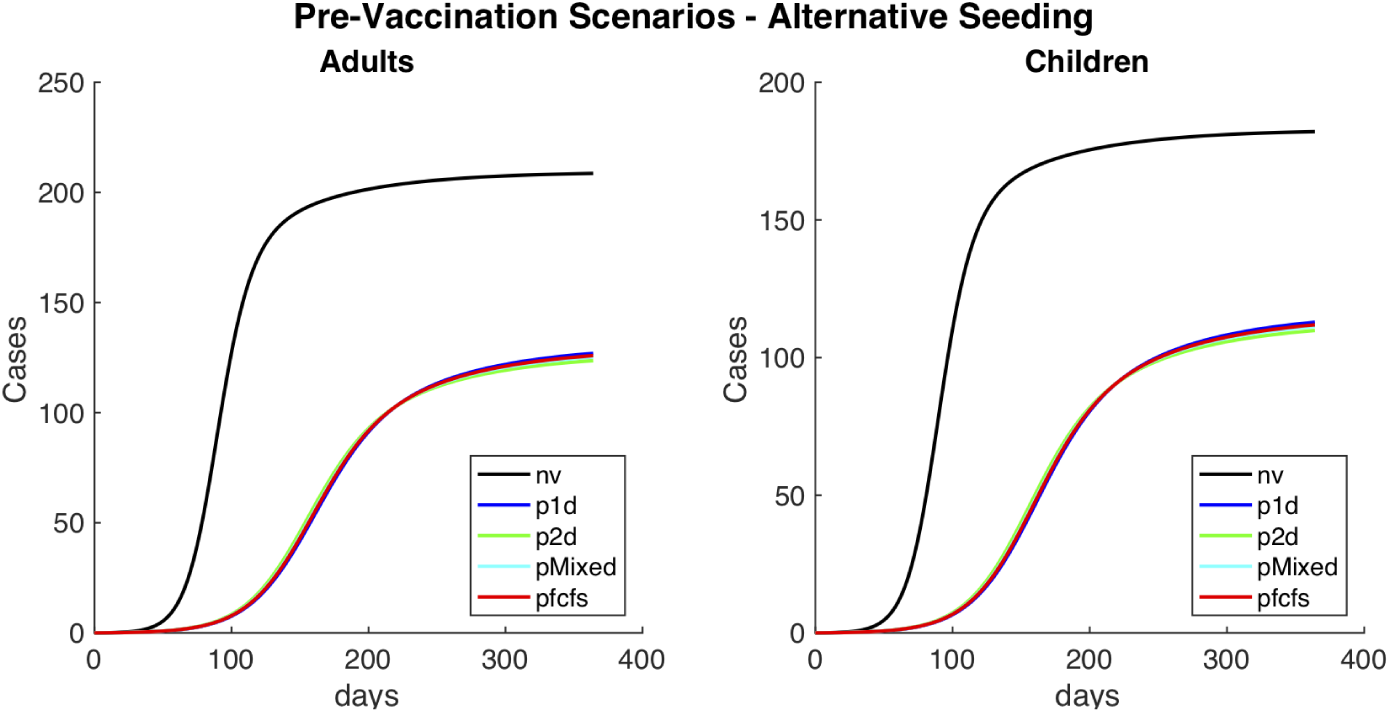
Alternative seeding scenario: one actual case. Cumulative cholera cases in adults and children, for different pre-vaccination scenarios. Where ‘nv’ is the baseline, no vaccination scenario; ‘p1d’ is the one-dose scenario; ‘p2d’ is the two-dose scenario; ‘pmixed’ is the mixed scenario.

**Figure 11.**
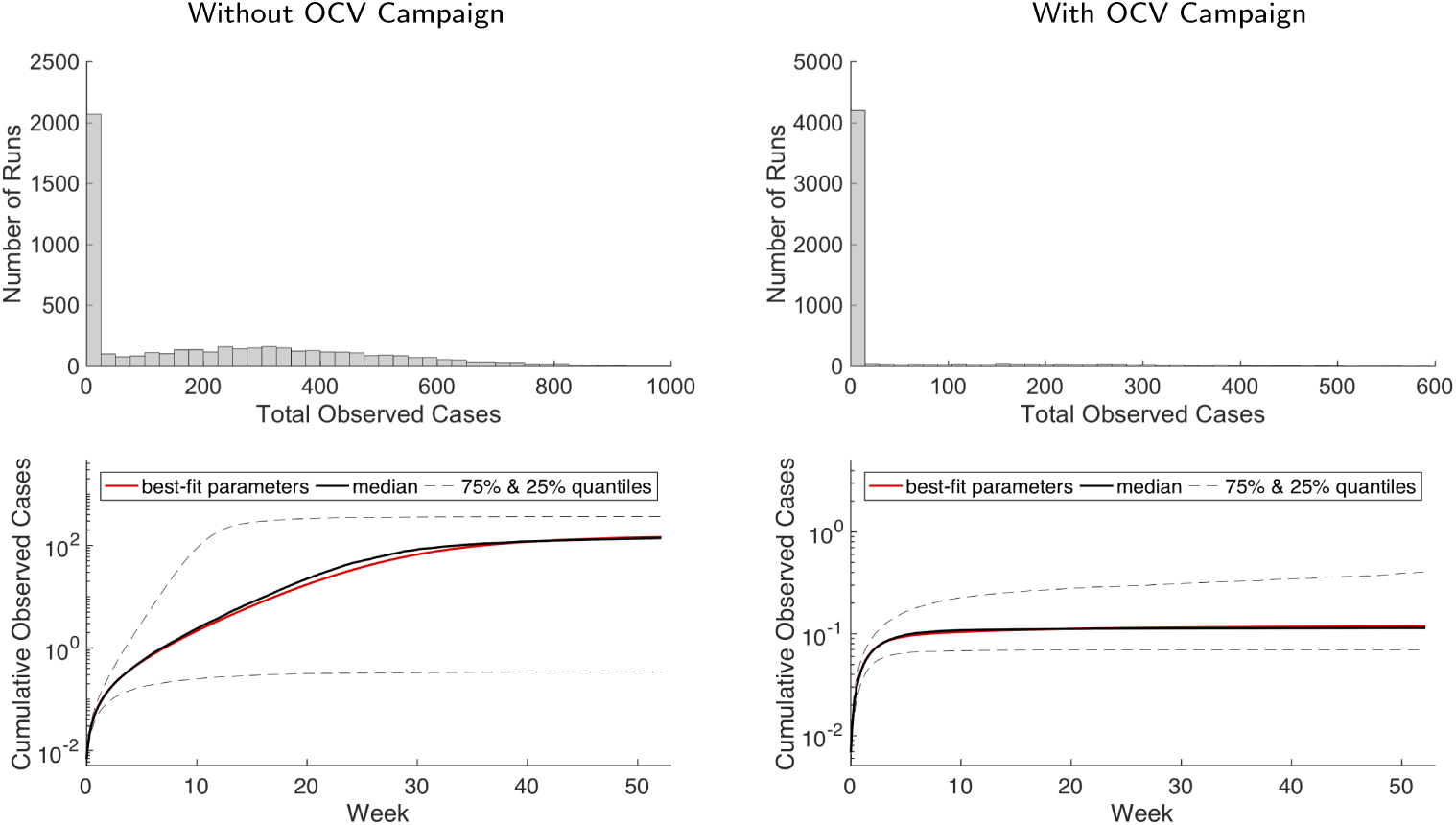
2013 forecast with a single actual case in adults and children as seeding. These plots show a partially immune population with no OCV campaign on left and with OCV campaign on right.

#### Forecasts for the 2014 Cholera Season

The 2014 forecasting results are quite similar to the 2013 runs. The partially immune population results in more simulations with 0 total cases, compared with the fully susceptible population. Because population immunity wanes between 2013 and 2014, there is higher proportion of larger outbreaks for the 2014 forecasting scenarios, but the vast majority of runs remain close to 0 for both populations. For details, see Supporting Information Figure 12.

**Figure 12.**
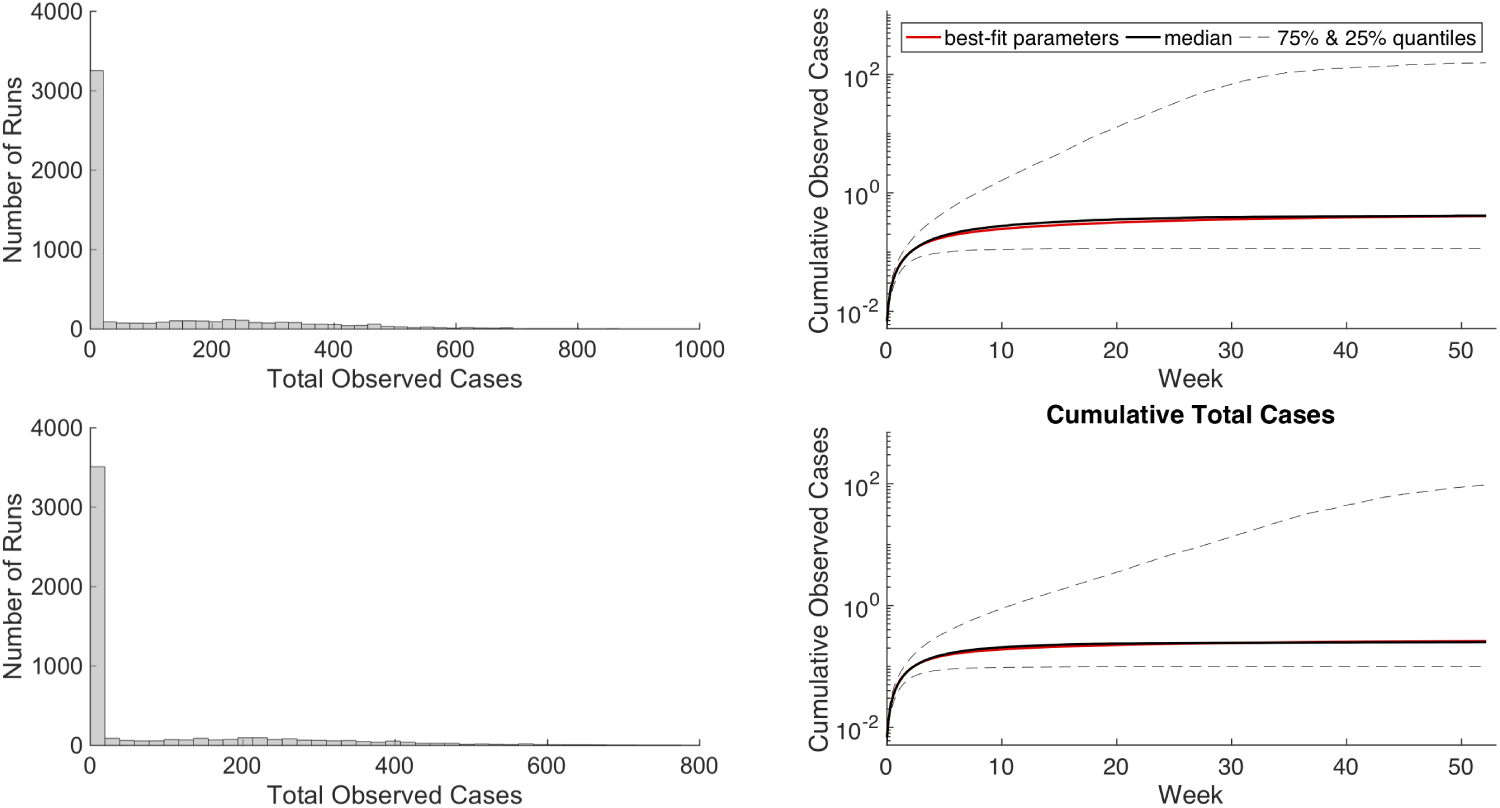
2014 forecast with a single actual case in adults and children as seeding. First row: Fully susceptible population with OCV campaign. Second row: Partially immune population with OCV campaign.

## Discussion

Using a mathematical model of cholera transmission in a refugee camp, we have shown the dramatic effect that vaccination can have on reducing the risk of cholera outbreaks in refugee settings. Our analyses suggest that pre-vaccination campaigns, even if not fully covering the whole camp population, can lead to substantial reductions in the number of observed cases in the event of an outbreak. Of course, the best pre-vaccination strategy depends on both one-dose effectiveness and the amount of doses available. If resources are limited and one-dose offers acceptable protection, single dose-based strategies would be preferred. But if there are enough vaccine doses available or the effectiveness of a single dose is questionable, then full vaccination strategies are better. We also found that reactive vaccination strategies can result in a moderate reduction in the number of infections. Further, the small differences in attack rates in our model results suggest that, at the best-fit parameters, vaccinating more individuals with at least one dose (even if the full vaccination sequence is not completed) may be nearly or more effective than vaccinating fewer individuals with a complete two-dose program assuming conservative vaccine effectiveness estimates. This consideration is particularly important if vaccination occurs later in the outbreak, where time is of the essence as the outbreak is already ongoing—the full two-dose sequence requires a delay between doses, hindering its effectiveness in reactive vaccination campaigns that begin later in an epidemic. Additionally, if logistical constraints limit the ability to follow up and provide patients with a second dose, a one-dose strategy may be preferable. However, the relative one vs. two-dose effectiveness, timing and number of doses available should all be considered. Finally, our projections of future outbreaks after the 2013 vaccination campaign in the Maela refugee camp suggest that vaccination may have prevented outbreaks in 2013 and 2014, as no cases were observed in either year, both in the model and during follow-up in the camp. This ability to consider counterfactual scenarios and generate projections highlights the potential of modeling to help guide real-time public health decision-making.

Our mathematical model of cholera transmission reproduces the dynamics observed in Maela’s 2010 cholera outbreak (Figure 2). We conducted a theoretical exploration of the dynamics of our model to examine different vaccination scenarios with only 20,000 doses of Shanchol distributed in the camp to reflect potential public health intervention strategies. Doses were distributed proportionally among children and adults. These results provide insight into the most effective strategies for vaccination when logistics might impede complete coverage of a population with one or two doses. We found that the two-dose pre-vaccination strategy was marginally the most effective with an attack rate of 5.5 cases per 1,000 people, but others provided comparable protection. On the other hand, the most effective reactive vaccination scenarios are the one- sand two-dose strategies with attack rates of 5.9 cases per 1,000 people (Table 3), while again others provided similar protection. All similarities between scenarios are the result of the conservative effectiveness estimates we used such that one-dose effectiveness is nearly exactly half of two-dose effectiveness.

For pre-vaccination, all scenarios can achieve large reductions in case counts. If one-dose effectiveness is low or uncertain, the two-dose scenario is preferable since it can guarantee larger or more certain reductions in case counts i.e., 44,000 doses (administered to 22,000 people) are sufficient to achieve *<* 50 cumulative cases. On the other hand, if one-dose effectiveness is only ∼ 50% that of two-dose effectiveness, 45,000 doses (administered to 45,000 people) are sufficient (see Figure 5 for details). In the reactive vaccination scenarios, two doses may be preferable if one-dose effectiveness is low or uncertain. Further, we see a crucial interplay between timing and vaccine effectiveness on transmission. Past a certain point in the outbreak, one-dose effectiveness does not substantially change the total case counts. For instance at day 50 in the one-dose scenario, the largest potential difference in cases is 65. (total cases: 395.3 and 333.2 for a one-dose effectiveness of 0% and 63%, respectively) while at day 20 the largest potential difference in cases is 202.2 (total cases: 395.5 and 202.3 for a one-dose effectiveness of 0% and 63%, respectively). Additionally, when one-dose effectiveness is sufficiently high, the one-dose strategy is preferable, with one-dose able to achieve the lowest case counts during an idealized situation in which there is no delay in reactive vaccination administration and the one-dose effectiveness is nearly equal to the two-dose effectiveness. Although this scenario may not be possible in real-world settings, it underscores the necessity of considering delays and relative one vs. two-dose effectiveness in reactive vaccination campaigns. For the one-dose scenario considered here, more individuals are given vaccine over an 8 day campaign resulting in more people having some protection early in the outbreak. On the other hand, in the two-dose scenario individuals who have been vaccinated have a higher level of protection, but achieving the same amount of population level protection will take a longer period of time i.e., two 4 day campaigns occurring 14 days apart (see Figure 7 for details).

The results of the forecasting analysis for 2013 show that regardless of population level immunity, the OCV campaign using a partially mixed strategy (coverage levels shown in Table 6) prevents a majority of outbreaks that might otherwise have occurred, shown in Figures 8 and 12. Additionally, even if we assume a fully susceptible population prior to the 2013 OCV campaign, the vast majority of post-OCV campaign runs in 2014 still result in no outbreak. Indeed, the median of total cases across all runs is *<* 1 (see Table 4 and Figure 12). This suggests that an introduction of cholera would likely not have resulted in a significant outbreak even given a conservative assumption about the population level of immunity. Thus, it was determined that there was no need for a booster campaign in Maela (and indeed there was no cholera outbreak that year).

**Table 6.** OCV coverage data from the 2013 vaccine campaign. The “% Coverage” column indicates the percent coverage for the entire population (all included and excluded subjects). The “Number of Individuals” column indicates the initial conditions used in the model, calculated from the coverage percentages.

| Class | % Coverage | Number of Individuals |
| --- | --- | --- |
| Non-vaccinated infectious adults - seeding ($I_a$) | 0% | 72 |
| Non-vaccinated adults ($S_a$) | 0% | 7,856 |
| Once-vaccinated adults ($V_a$) | 22.3% | 6,208 |
| Twice-vaccinated adults ($VV_a$) | 49.3% | 13,765 |
| Non-vaccinated infectious children - seeding ($I_c$) | 0% | 53 |
| Non-vaccinated children ($S_c$) | 0% | 3,021 |
| Once-vaccinated children ($V_c$) | 18.7% | 3,241 |
| Twice-vaccinated children ($VV_c$) | 63.6% | 11,018 |

A key limitation of our model is the uncertainty in parameter values. To assess the uncertainty in our parameter values, we conducted global sensitivity analyses using LHS and assessed both practical and structural identifiability of the model. The quantitative results are heavily dependant on the vaccine effectiveness estimates however, we chose the lower bounds of values from recent studies to ensure that the model generated conservative results. Furthermore, the qualitative conclusions i.e. the interplay between vaccine timing and vaccine effectiveness will occur for most realistic effectiveness estimates. Other weaknesses of this analysis are inherent in the model assumptions. For example, one key assumption for the vaccination scenario simulations is that there is no waning immunity because we are simulating over a short time course. We do, however, incorporate waning immunity into the forecasting analysis. Another assumption is that each infectious individual is equally infectious regardless of previous vaccination, cholera exposure, or time since infection. In any case, this assumption would result in an over-estimate of the total number of cases because we are ignoring the fact that individuals who have been vaccinated might be less infectious. We are also ignoring hyperinfectiousness, as incorporating this into our model as including it would require tracking of pathogen through the human host and a shorter time scale because it decays after 18 hours [76]. Although we are not explicitly modeling asymptomatic infections or errors in disease reporting, these are accounted for in the scaling factors, *k*_*a*_ for adults and *k*_*c*_ for children. Additionally, in our analyses we used a deterministic model which will not capture stochastic fluctuations. Stochasticity may play a role, particularly in the early phase of an outbreak following a new introduction of cholera—our model may therefore miss some of the stochastic die-out of epidemics in our forecasts, which again, would lead to an overestimate of the number of cases. We assume that the mortality rate for individuals with cholera is the same as that for individuals without cholera. Although this is likely not true, the data we fit our model to did not have a sufficient number of deaths to calculate case fatality rates for diseased compared with non-diseased population groups (there was only one death among individuals with cholera). As individuals in Maela have easy access to medical care, they are likely to be treated and have a high rate of survival. For the sake of parsimony [77], we are also assuming that human-human transmission parameters are equal across demographic groups and separately human-water transmission parameters are equal since we obtained similar fits for a range of *β* values in the practical identifiability analysis (see Supporting Information). We are using data from a specific setting, which may limit the external validity of our results; however, the qualitative dynamic results obtained should be consistent for other refugee camp settings and have been seen in similar mathematical modeling analyses [43, 78].

The strengths of this model include the novelty of considering vaccination scenarios while explicitly accounting for environmental transmission of cholera in a refugee camp. Additionally, our identifiability and sensitivity analyses methodically considered parameter uncertainty. Another strength is the fact that we used real-world data to inform our model which in turn, provided insight for the public health response. Specifically, we liaised with the CDC as well as local nongovernmental organizations in real-time and used data from the OCV campaign to directly inform the model. Further, the model was then used to explore counterfactual forecasting scenarios to help answer outstanding questions among trial investigators about whether or not a booster campaign was necessary. Finally, the consistency of our results with other analyses indicates the robustness of our findings.

In general, our results indicate that vaccination should be considered in conjunction with WaSH with the caveat that immunity may wane over time. Furthermore, the trade-off between vaccine effectiveness (i.e., one-dose compared with two-dose) and timing of reactive vaccination should be carefully considered. The WHO currently holds a stockpile of over 3 million doses of oral cholera vaccine to allow countries or institutions to request doses of vaccine during cholera outbreaks. The average time from when requests were approved to receipt in country has been 14.4 days with an additional days until vaccination actually started [79, 80]. As seen in our vaccination scenario results, timing is crucial to the impact of vaccination. Recent mathematical modeling work has examined how best to allocate global stockpile reserves [81].

Ideally, pre-vaccination should be considered as a short-term transmission reduction strategy, compared to the potentially longer-lasting effects of improved WaSH. A recent study used a static model fit to data from Malawi to estimate cases averted in Haiti by implementation of oral cholera vaccine and/or WaSH and found that a combined implementation of WaSH and vaccine resulted in the greatest reductions in cases [82]. Furthermore, WaSH reduces transmission for a wide range of infectious diseases and if maintained is more permanent, while Shanchol targets cholera, and its effects do not last as long—although vaccine campaigns may be easier to implement, as they do not require sustained maintenance. Thus, both WaSH and vaccination may have their roles to play in an effective intervention strategy.

Our analyses suggest that vaccination provides an effective strategy for preventing cholera outbreaks in refugee camps and that cholera vaccination should be considered, even in the absence of an ongoing outbreaks. Given the dramatic increases in displaced populations and refugee settlements across the world it is critical that vaccination be considered with water sanitation and hygiene improvements. If a camp is facing an outbreak, delayed distribution of vaccines can substantially alter the effectiveness of a reactive vaccine campaign, suggesting that quick distribution of vaccines (e.g., using a first come, first served approach) may be more important than ensuring that every individual gets both vaccine doses.

## Conclusions

We developed an age-structured SIWR-based transmission model to consider different cholera vaccination strategies in Maela, the largest and most long-standing refugee camp in Thailand. Our model was fit to cholera incidence data from 2010 and was parameterized using demographic data collected from the camp. We considered multiple scenarios, including both a theoretical exploration of the effects of variation in timing, effectiveness and supply, as well as the real-world coverage of vaccine in Maela. We found that the preferred number of doses per person and timing of vaccination campaigns should be considered in the context of one vs. two dose effectiveness and logistical constraints. Importantly, our analysis coincided with an actual cholera vaccination campaign in the camp and was used to evaluate the campaign and to help determine that there was no need for a follow-up booster campaign. The setting of our analysis is particularly relevant given the recent worldwide increase in total numbers of refugees. Results from our model highlight the utility of vaccination to prevent cholera. Vaccination campaigns can be combined with more permanent water, sanitation, and hygiene infrastructure improvements to reduce the risk of cholera and other enteric disease epidemics. Overall, this study demonstrates that mathematical modeling can generate useful insights into real-time intervention decisions.

## List of Abbreviations

(WaSH): Water, Sanitation and Hygiene (WaSH)
(OCV): Oral Cholera Vaccine
(WHO): World Health Organization (WHO)
(CDC): Centers for Disease Control and Prevention
(SIWR): Susceptible-Infectious-Water-Recovered model
(PU-AMI): Premiére Urgence Aide Medicale Internationale (PU-AMI)
(VE): Vaccination Effectiveness (VE)

## Declarations

### Ethics approval and consent to participate

This study was determined to be nonregulated by the University of Michigan institutional review board (HUM00061865) because only deidentified data were used. Therefore, the need for consent was not required for this study, however in the original data collection (see [15] for more details), entry screeners obtained consent verbally (as illiteracy is high). Deidentified data was shared with the authors as part of an ongoing collaboration between JH, RM, and MCE with the CDC and its partners. However, CDC and its partners retain ownership of the data and approved its use in publication.

### Consent for publication Not Applicable

### Availability of data and material

The datasets used and/or analysed during the current study are available from the corresponding author on reasonable request.

### Authors’ contributions

JH, RM and MCE came up with the research plan, analyzed the data, and wrote the paper. CRP provided guidance on the dataset and KD and CRP provided guidance on the analysis plan and paper. All authors provided edits and approved the final manuscript.

## Acknowledgements

We would like to thank Dr. Nuttapong Wongjindanon Thailand Ministry of Public Health – U.S. Centers for Disease Control and Prevention Collaboration, Nonthaburi, Thailand for his helpful comments, advice, and support. Furthermore, we would like to thank PU-AMI and Niamh de Loughry in Maesot, Thailand, for providing us with key data and information for our analyses and for helping us gain access to and navigate Maela.

## Supporting Information

**1 Profile Likelihoods for Fitted Parameters**

**2 Cumulative Cases of Alternative Seeding Scenarios**

**3 Cumulative Cases of Alternative Seeding Scenarios**

**4 Maela Vaccination Coverage Calculations**

**5 2013 Forecasting Results with Partially Immune Population**

**6 2014 Forecasting Results**

### 7 Model equations and additional details

#### 7.1 Simplified Model Equations

The simplified age-structured model equations with no vaccination are below. This model was used for the identifiability analysis to calculate ℛ_0_ and was fit to the Maela outbreak data.

*Force of Infection Equations*

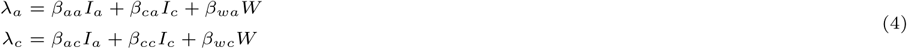

*Non-Vaccinated Adults*

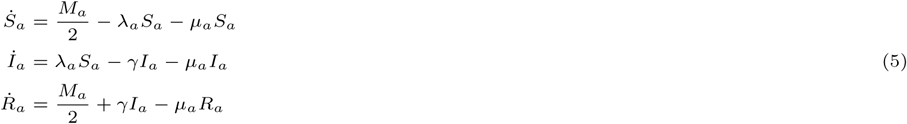

*Non-Vaccinated Children*

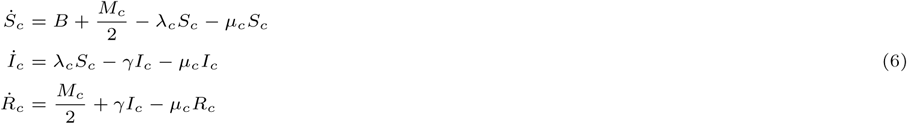

*Environmental Pathogen*

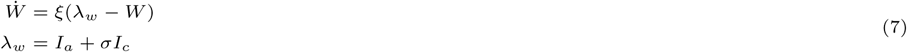

#### 7.2 Full Model Equations

The full age-structured model equations separated by non-vaccinated, once-vaccinated, and twice-vaccinated individuals are below. If fitted, parameter values are from the simplified model (above) and the remaining non-fitted values (e.g., vaccine effectiveness) are from the literature, see Table 1. This model was used to examine the different counterfactual vaccination scenarios.

*Force of Infection Equations*

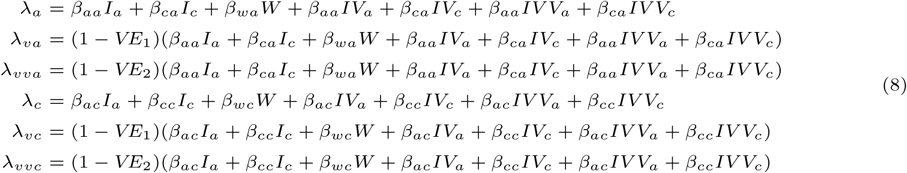

*Non-Vaccinated Adults*

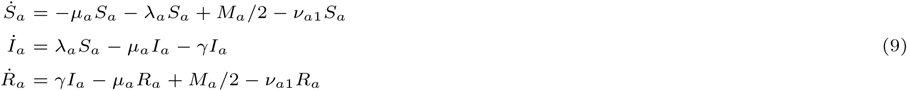

*Once-Vaccinated Adults*

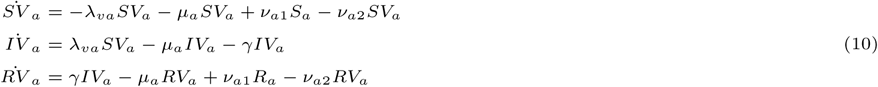

*Twice-Vaccinated Adults*

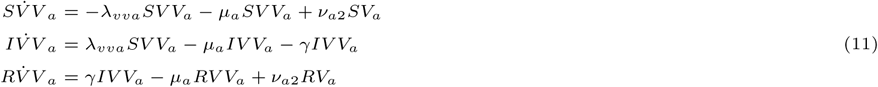

*Non-Vaccinated Children*

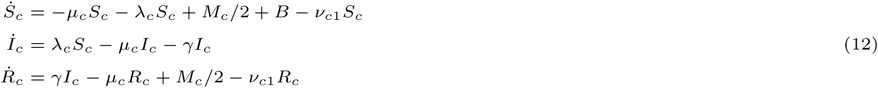

*Once-Vaccinated Children*

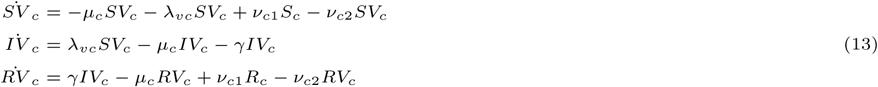

*Twice-Vaccinated Children*

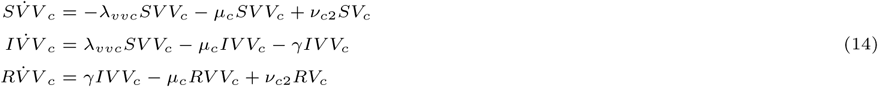

*Environmental Pathogen*

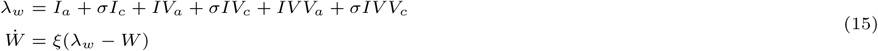

*Total Population Sizes*

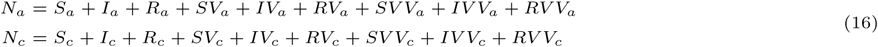

#### 7.3 Forecasting Model equations

The age-structured model equations used for the forecasting scenarios are below.

*Non-Vaccinated Adults*

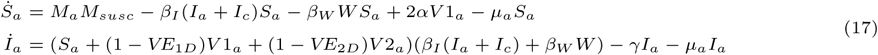

*Non-Vaccinated Children*

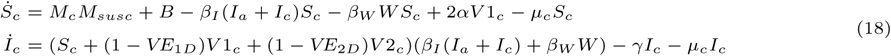

*Immune Adults*

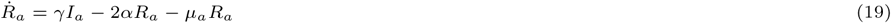

*Immune Children*

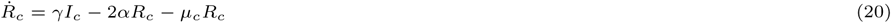

*Partial Immune/Vaccinated Adults*

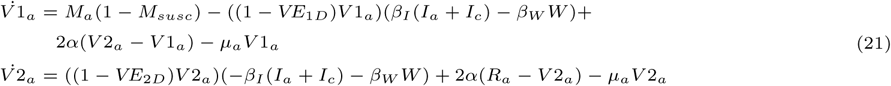

*Partial Immune/Vaccinated Children*

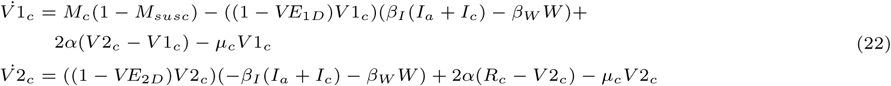

*Environmental Pathogen*

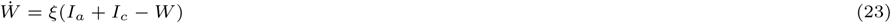

##### Identifiability Analysis

Identifiability analysis addresses the question of whether the model parameters can be estimated from a given data set [30]. Identifiability is typically broken into two broad categories—(1) structural identifiability, which examines theoretical identifiability from the structure of the model and measured variables, and (2) practical identifiability, which addresses how a model’s identifiability properties are affected by real-world data issues such as noise and sampling frequency.

#### Structural Identifiability Analysis

To examine the structural identifiability of our simplified age-structured model, we used the differential-algebra based approach developed in [54, 30, 55, 56, 57, 58]. Determining the structural identifiability of the model is a prerequisite to determining if there is a unique solution for a set of unknown model parameters [54]. Structural identifiability can be framed as evaluating whether the model parameters can be estimated uniquely, when the data is assumed to be ‘perfect’ (i.e., noise-free and measured for all time points). Establishing structural identifiability is a prerequisite for successful parameter estimation from real-world, noisy data. When parameters are not individually identifiable, groups of parameters typically form identifiable combinations that can be uniquely determined.

In the differential algebra approach, the unmeasured state variables (e.g. *S*_*A*_, *S*_*C*_, etc.) are eliminated, leaving equations only the measured variables, their derivatives, and the parameters, denoted the input-output equations. In this case, the measured variables are cholera incidence among adults and children. The identifiability from cholera incidence was more easily analyzed using the prevalence approximation, which as *γ* is assumed to be known, yields the same structural identifiability results as the standard incidence. We assumed the demographic parameters, initial population sizes, and recovery rate are known from data as described above and defined in Table 1, and the remaining parameters (*β*_*ij*_’s, *k*’s, *α, σ*, and *ξ*) were considered unknown. A Gröbner-basis approach was then used to test whether the unknown model parameters in Equations (4) – (7) are identifiable from the measured data, with all calculations performed in Mathematica Version 10.

Similar to the original SIWR model [30], the waterborne transmission parameters and *α* were not separately identifiable for our model, instead forming the identifiable combination 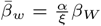*>*. To address this, we define 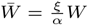. Rewriting the model equations in terms of these new variables yields the following equation for environmental pathogen:

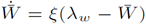

with all other equations remaining the same except replacing *W* with 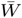 and the identifiable combination 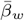. Once re-scaled, all unknown model parameters 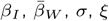, *σ, ξ*, and the *k*’s) were structurally identifiable. From this point forward (and for the parameter estimation and other analyses), we use only the rescaled versions of *β*_*W*_ and *W*, and thus we will omit the bar notation.

#### Practical Identifiability

Initially, even though structural identifiability was considered, we obtained extremely similar fits for a wide range of transmission parameter values, suggesting that there were practical unidentifiability issues wherein the reporting parameters (*k*_*a*_ and *k*_*c*_) and adult and child transmission parameters can partially compensate for one another to yield the same overall apparent cholera incidences. For the sake of parsimony [77], we set all human-human transmission parameters equal to each other, denoted *β*_*I*_, and separately we set all human-water transmission parameters equal to each other, denoted *β*_*W*_. Similarly, *σ*, the relative shedding rate for adults and children, was also relatively practically unidentifiable, and so we set shedding to be equal for both classes.

To examine practical identifiability and parameter uncertainty, we plotted profile likelihoods of each fitted parameter. Profile likelihoods are a numerical approach to evaluating parameter uncertainty and identifiability [59]. Profiles are generated by fixing the profiled parameter to a series of values, while fitting the remaining parameters that are being estimated. Typically, the minimum negative log likelihood (or equivalently the maximum likelihood) values are plotted for each value of the profiled parameter, forming the profile likelihood for that parameter. The minimum represents the best-fit value of the profiled parameter and is determined by parameter estimation. If the profile is flat, the parameter cannot be uniquely determined and is considered unidentifiable. However, even if the profile is structurally identifiable, the curvature may be quite shallow, so that a particular minimum cannot practically be distinguished - this is denoted practical unidentifiability. Confidence intervals can be determined from the profile likelihood by setting a significance-based threshold on the likelihood based on a *χ*^2^ distribution [59]. Once the threshold is set, all parameters corresponding to likelihood values below the threshold fall within the confidence interval. The results from the profile likelihood plotting can be seen in Figure 9.

##### Sensitivity Analysis: Initial Seeding from Observed to Actual

As another sensitivity analysis, we changed our initial seeding from one observed case to one actual case for the Maela and pre-vaccination scenarios. Because vaccination occurred before any outbreak, administration of the vaccine was not affected by case detection. Overall, we see the same pattern of results, shown in Figure 10 and Table 5. The two-dose scenario sees the largest reduction in cases followed by the mixed, first come, first served, and one-dose scenarios. Since the number of initial infected individuals is lower, the total cumulative case counts are as well. Of note is that the reduction in cases is greater for pre-vaccination scenarios than for the baseline non-vaccination scenario.

##### Maela Vaccine Coverage Calculations

Among included individuals (pregnant women and infants *<* 1 year were excluded), the OCV campaign covered 51% of adults and 68% of children with two doses, and another 23% of adults and 20% of children with one dose. We made the following adjustments to determine total Maela coverage among both included and excluded individuals for the forecasting scenarios:

- Once-vaccinated adults: (*V*_*a*_ *-V V*_*a*_)((*N*_*a*_*-*Pregnant women)*/*(*N*_*a*_))*N*_*a*_ ((0.74-0.51)*(27901-910)/27901)*27901 = **6207.9**
- Twice-vaccinated adults: (*V V*_*a*_)((*N*_*a*_*-*Pregnant women)*/*(*N*_*a*_))*N*_*a*_ (0.51*(27901-910)/27901)*27901 = **13765.4**
- Once-vaccinated children: (*V*_*c*_ *-V V*_*c*_)((*N*_*c*_*-*infants under 1 year old)*/*(*N*_*c*_))*N*_*c*_ ((0.88-0.68)*(17332-1129)/17332)*17332 = **3240.6**
- Twice-vaccinated adults: (*V V*_*c*_)((*N*_*c*_*-*infants under 1 year old)*/*(*N*_*c*_))*N*_*c*_ (0.68*(17332-1129)/17332)*17332 = **11018**

Table 6 shows the total number of individuals by class used to simulate the OCV campaign.

##### Forecasts for the 2013 Cholera Season

In the 2013 forecasting results, we see a larger spread of total case numbers for runs in the scenario without the OCV campaign compared to the scenario with the OCV campaign. The partially immune population runs generally have lower case counts when comparing to the fully susceptible population. Furthermore, for the scenarios that consider the OCV campaign, we see that the vast majority of runs having case counts close to 0. For details see Figures 8 and 11.

##### Forecasts for the 2014 Cholera Season

The 2014 forecasting results are quite similar to the 2013 runs for the fully susceptible population compared to the partially immune population with more runs resulting in 0 total cases for the partially immune population. As population immunity wanes between 2013 and 2014 we get a higher proportion of larger outbreaks for the 2014 forecasting scenarios, but the vast majority of runs remain close to 0 for both the fully susceptible and partially immune populations. For details see Figure 12.

